# Biology-guided machine learning for robust anatomical correspondence across biodiversity datasets

**DOI:** 10.64898/2026.09.28.755222

**Authors:** Arthur Porto, Katherine A. Wolcott, Kelsey L. Fenner, Caleb Gordon, Breanna Shi, Edward L. Stanley

## Abstract

Three-dimensional imaging is rapidly increasing the scale of biological morphometric datasets, but establishing automated anatomical correspondence remains a major bottleneck. Conventional point-cloud registration methods transfer landmarks efficiently, yet poor initialization or high morphological variation can produce geometrically plausible but biologically implausible deformations.

Here, we developed a statistical-shape-model (SSM)-regularized coherent point drift (CPD) estimator for automated morphometric landmark transfer. It replaces the unconstrained non-rigid CPD update with optimization in SSM coefficient space and regularizes coefficients by inverse population covariance, so deformations along low-variance biological modes are penalized more strongly. We compared pose-matched conventional CPD, SSM-CPD, and SSM-CPD with refinement in leave-one-out atlas experiments on laboratory mouse skulls (N=249), primate molars (N=115), and squamate vertebrae (N=254) under aligned, mildly perturbed, and severely perturbed initialization. We evaluated these formulations in terms of landmark accuracy, failure-tail behavior, runtime, and preservation of morphometric structure. Under severe perturbation, we also evaluated an integrated recovery procedure that optimizes the starting template, recovers global pose, and then applies SSM-CPD with refinement.

Severe misalignment produced widespread failures in pose-matched conventional CPD, whereas SSM regularization reduced median error from 13–21% to approximately 1–3% of the target bounding-box diagonal. This improvement was highly consistent for skulls and molars. In the morphologically complex vertebral dataset, however, low median error concealed a substantial minority of catastrophic failures (28% of vertebrae above a 6% error threshold), which integrated recovery reduced to 4%. Across datasets, automated landmarks preserved the major axes and pairwise relationships of morphometric shape space, and local CPD refinement improved covariance preservation.

Embedding learned biological variation directly into registration broadens the range of starting conditions and anatomies for which automated landmark transfer remains reliable. Our results also show that failure tails and downstream morphometric fidelity must be assessed alongside median accuracy when automated landmarking methods are evaluated.

## 1 Introduction

Quantitative analyses of organismal form depend on anatomical correspondence: the assignment of comparable biological meaning to points, curves, surfaces or regions across specimens. In geometric morphometrics, this correspondence is commonly represented by homologous landmarks and semilandmarks, which provide the coordinate system through which disparity, allometry, modularity, integration, and evolutionary rates are estimated (Bookstein, 1991; Rohlf & Slice, 1990; Dryden & Mardia, 1998; Mitteroecker & Gunz, 2009; Bardua *et al*., 2019). Because correspondence defines the variables that enter subsequent analyses, inconsistent landmarking can cause measurement error to be interpreted as biologically meaningful (Fruciano, 2016; Mitteroecker & Schaefer, 2022).

The growth of 3D biological imaging has shifted the bottleneck in morphometrics from image acquisition to annotation. Museum digitization, computed tomography, surface scanning, photogrammetry, and increasingly efficient segmentation workflows now make it possible to assemble surface models at unprecedented scales (Hedrick *et al*., 2020; Blackburn *et al*., 2024; Hsiang *et al*., 2018; Lürig *et al*., 2021). However, producing reliable landmarks remains labor-intensive, especially when datasets span large morphological disparity (Percival *et al*., 2019; Porto & Voje, 2020). Automated landmark transfer offers a practical route to scalable morphometrics, but only if registration remains stable when shapes are variable and initialization is imperfect.

Volumetric single- and multi-atlas registration provided an important early route to automated landmark transfer in mouse morphometrics (Young & Maga, 2015; Maga *et al*., 2017; Percival *et al*., 2019; Devine *et al*., 2020). SlicerMorph’s ALPACA subsequently helped make surface point-cloud landmark transfer practical for three-dimensional biological morphometrics by placing point-cloud registration within an accessible 3D Slicer workflow (Fedorov *et al*., 2012; Rolfe *et al*., 2021; Porto *et al*., 2021). Subsequent multiple-template approaches showed that template-based landmarking can be extended to more variable samples by selecting among alternative source specimens (Zhang *et al*., 2022). Parallel work has developed landmark-free surface metrics, globally transitive pseudolandmark correspondences, and data-driven landmark sampling for 3D morphometrics (Boyer *et al*., 2011, 2015; Gao *et al*., 2018, 2019; Koehl & Hass, 2015). Together, these approaches expand sample size and surface coverage, while making correspondence accuracy and consistency key methodological concerns.

Template-transfer methods nevertheless share a limitation with most point-cloud registration algorithms: the correspondence problem is solved from geometry alone. Coherent point drift (CPD), which is used by both ALPACA, treats one point set as Gaussian-mixture centroids and estimates a transformation that maximizes correspondence likelihood while regularizing motion coherence (Myronenko & Song, 2010). This probabilistic formulation is attractive for biological surfaces because it is smooth and relatively robust to noise. Nevertheless, a purely geometric deformation can fail when initial alignment is poor or when local geometric similarity is ambiguous. In such cases, the algorithm may find a low-distance solution that is geometrically plausible but anatomically incorrect.

What these methods lack is biological information about which deformations are plausible. In other words, their deformation penalties are not informed by observed population covariance. Anatomical variation is structured by development, function, and phylogeny, so empirical shape covariance contains information about plausible deformation (Klingenberg, 2008; Porto *et al*., 2009; Melo *et al*., 2016; Mitteroecker & Schaefer, 2022). Statistical shape models provide a direct way to represent this structure by encoding a mean configuration, principal directions of variation, and their corresponding variances (Cootes *et al*., 1995; Heimann & Meinzer, 2009).

Building on the broader principle that learned shape structure can constrain probabilistic registration (Cootes *et al*., 1995; Heimann & Meinzer, 2009; Hirose, 2017), we develop and openly implement a biology-guided CPD framework for automated morphometric landmark transfer. The SSM-CPD estimator developed here replaces the unconstrained non-rigid CPD parameters with optimization in SSM coefficient space, uses population eigenvalues to define covariance-weighted regularization, and jointly estimates similarity pose. An optional conventional CPD stage restores local geometric variation after the constrained fit. We also develop an integrated recovery estimator that searches the learned shape space for a target-compatible template, recovers global pose, and then applies SSM-CPD with local refinement.

We benchmark these implementations across three anatomically distinct datasets (laboratory mouse skulls, primate molars, and squamate vertebrae) under matched joint-similarity optimization. Using leave-one-out atlas landmarking, we quantify sparse landmark accuracy, robustness to poor initialization, and preservation of downstream morphometric structure. A focused factorial analysis separately evaluates the effects of pose optimization and registration formulation, and the severe-perturbation analysis evaluates the integrated recovery estimator. Together, these tests ask whether constraining registration by empirically observed biological shape variation can make automated landmark transfer more robust, anatomically plausible, and useful for downstream comparative morphometrics. The SSM-CPD estimator is distributed in the open-source biocpd library, and the complete registration and recovery workflows are integrated into MorphoWeave, a 3D Slicer extension for atlas construction and morphometric landmark transfer.

## 2 Materials and Methods

### 2.1 Study design overview

We tested whether constraining CPD with a statistical shape prior improves automated landmark placement relative to conventional CPD under equivalent joint similarity optimization, particularly when target specimens are initially misaligned (Fig. 1). The benchmark used a leave-one-out atlas design across three anatomical datasets. For each held-out target, we built a statistical shape model from the remaining specimens, registered the dense atlas correspondences to the target surface, transferred the sparse atlas landmarks through the resulting deformation, and compared predicted landmarks with expert annotations (Table 1). We evaluated performance across aligned, mildly perturbed, and severely perturbed initialization regimes. Under severe perturbation, we separately evaluated the integrated recovery estimator.

**Figure 1:**
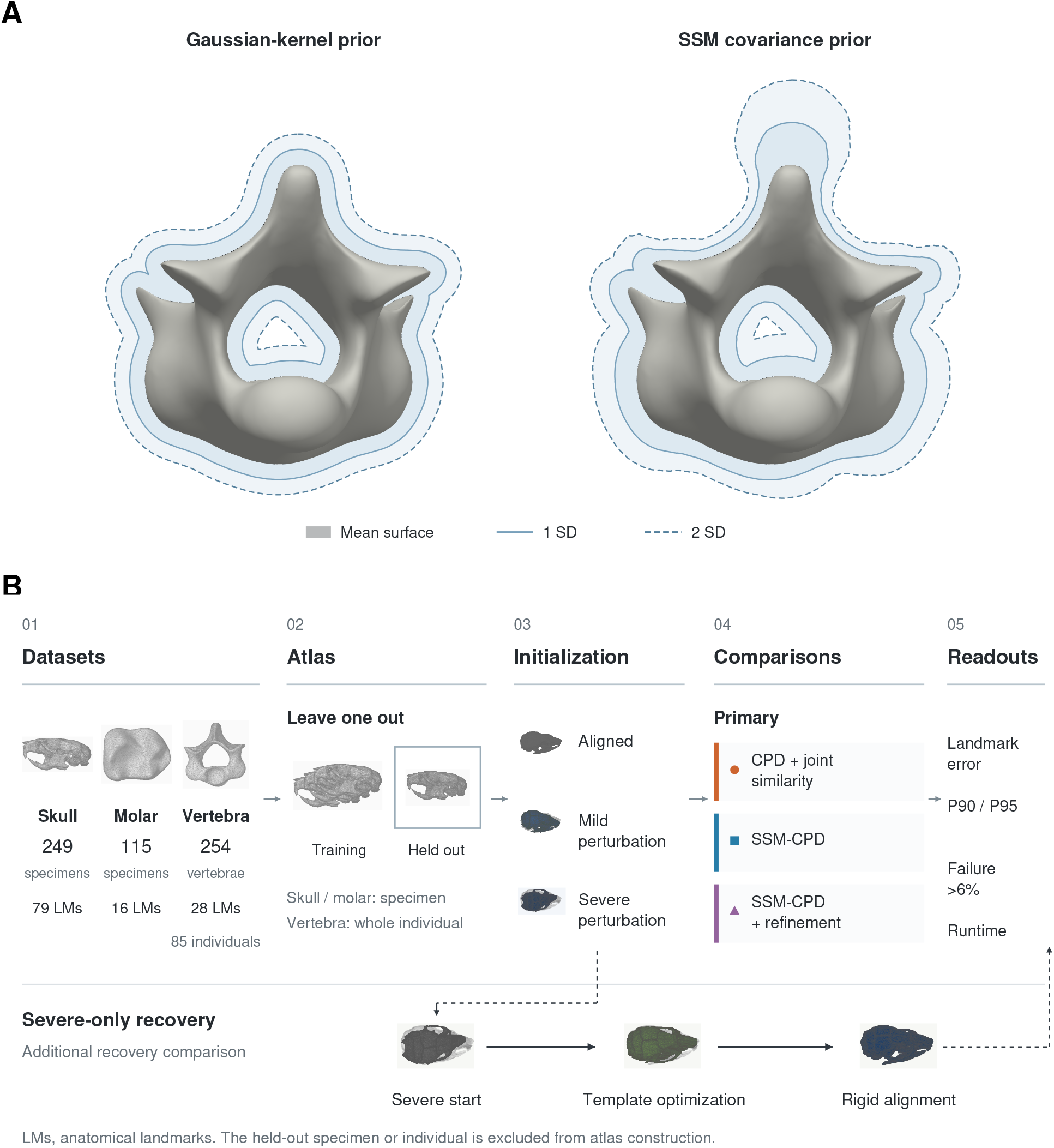
Method concept and experimental design. **(A)** Schematic contrast between the deformation envelope implied by a Gaussian-kernel CPD prior and that implied by the SSM covariance prior. The grey surface is the mean vertebra; solid and dashed outlines mark 1 and 2 standard deviations of plausible deformation. Envelopes are illustrative and are not fitted parameter maps. **(B)** Datasets, leave-one-out construction (whole individuals held out for vertebrae), aligned, mild, and severe initialization regimes, the three pose-matched primary estimators, the severe-only recovery comparison, and the evaluation readouts.

**Table 1:** Dataset size, landmark dimensionality, dense-correspondence count, and leave-one-out inference unit.

| Dataset | Specimens | Sparse LM | Dense points | LOO unit | Anatomical challenge |
| --- | --- | --- | --- | --- | --- |
| Mouse skull | 249 | 79 | 2,665 | Specimen | High landmark count; limited disparity |
| Molar | 115 | 16 | 2,621 | Specimen | Compact structure; local geometric ambiguity |
| Squamate vertebra | 254 | 28 | 3,751 | Individual (85) | High disparity; repeated serial anatomy |

LM, landmarks; LOO, leave one out. Vertebral folds exclude every vertebra from the held-out biological individual.

### 2.2 Benchmark datasets and atlas preparation

The benchmark used three datasets representing different anatomical structures and landmarking problems: an unpublished laboratory-mouse skull dataset, primate molar surfaces derived from published anatomical-surface resources (Boyer *et al*., 2011), and squamate pre-sacral vertebral surfaces extracted from museum computed-tomography data made accessible through the openVertebrate initiative (Blackburn *et al*., 2024) (Table 1). The mouse sample represents a single model-organism system, whereas the molar and vertebral samples span broader taxonomic and morphological variation. The molar sample contains 115 second mandibular molars representing 45 primate species-level taxa and six close-relative outgroup taxa. The vertebral sample contains 84 squamate individuals spanning approximately 80 taxa across more than 30 families, together with one tuatara (*Sphenodon punctatus*), that serves as an outgroup. Each dataset contains aligned surface models, expert-applied sparse landmarks, dense population correspondences, and an atlas surface model with sparse and dense correspondences. The dense correspondences were generated using the DeCA dense correspondence framework of Rolfe & Maga (2023) as implemented in MorphoWeave. DeCA combines homologous landmarks with dense surface registration to create anatomically interpretable surface correspondence across specimens. These dense correspondences were used to construct the leave-one-out SSMs and the statistical atlas. Manually annotated sparse expert landmarks served as the reference standard for evaluating automated landmark transfer.

The mouse and molar datasets contain one specimen per biological unit, so each leave-one-out fold excludes one specimen. The vertebral dataset contains 254 vertebrae from 85 individuals. In this dataset, “specimen” refers to a single vertebra. Each held-out vertebral fold therefore excluded all vertebrae from the same individual, preventing within-individual leakage into the training SSM.

### 2.3 Leave-one-out SSM construction

For each fold, dense correspondence configurations from the training set were vectorized and used to build a statistical shape model by PCA (Cootes *et al*., 1995; Heimann & Meinzer, 2009). The SSM retained enough components to explain 99% of the variance in the dense correspondences. For registration, the model was further truncated to the subspace explaining 95% of the training-set variance. This distinction reflects the different goals of atlas construction and registration: the full atlas database preserves population-level variation, whereas the registration estimator benefits from a more conservative subspace that reduces overfitting to local geometric noise.

Let **Y**_0_ ∈ ℝ*^M^*^×^*^D^* denote the base atlas correspondence configuration and let **y**_0_ = vec(**Y**_0_). Let **U** ∈ ℝ*^MD^*^×^*^K^* contain the retained SSM basis vectors and let **Λ** = diag(*λ*_1_, . . . , *λ_K_*) contain the corresponding eigenvalues. The SSM deformation is parameterized by coefficients **b** ∈ ℝ*^K^*:

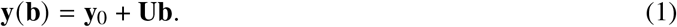

The deformed configuration **Y**(**b**) is obtained by reshaping **y**(**b**) into an *M* × *D* matrix. Thus, the non-rigid component is expressed within the biological variation observed in the training sample rather than as an arbitrary displacement field. The full estimator derivation is provided in Supplementary Note S1.

### 2.4 Initialization regimes

The primary robustness benchmark evaluated three initialization regimes. The aligned regime used the target meshes as provided after rigid registration. The mild perturbation regime applied seeded pseudorandom similarity perturbations whose three Euler angles (about the *x*, *y*, and *z* axes) are each drawn uniformly from ±15 degrees (composite rotation angle median 15 degrees, range 2–26 degrees), with translations drawn uniformly within ±5% of the target bounding-box diagonal along each axis and scale sampled from 0.9 to 1.1. The severe perturbation regime drew each Euler angle with a magnitude of 30 to 45 degrees and a random sign (composite rotation angle median 61 degrees, range 49–83 degrees), with the same translation and scale distributions. Each perturbation was seeded by a global seed combined with the specimen identifier, so every method received the same perturbed target. When a target was perturbed, the target mesh and reference landmarks were transformed together so that evaluation remains in the target coordinate system.

### 2.5 Primary registration methods

The primary comparison included three estimators with equivalent joint similarity optimization (Table 2), and we use the labels defined there throughout. CPD + joint similarity jointly estimates rotation, translation, scale, and a conventional deformable CPD field without an SSM prior (Myronenko & Song, 2010). SSM-CPD is the estimator developed for this study and implemented in biocpd. Its M-step updates SSM coefficients under an eigenvalue-weighted prior while jointly optimizing similarity pose. SSM-CPD + refinement first runs the complete SSM-CPD estimator and then applies conventional deformable CPD to refine the local geometric fit.

**Table 2:** Registration and recovery methods used in the robustness and severe-perturbation analyses.

| Method | Procedure | Primary role |
| --- | --- | --- |
| CPD + joint similarity | Coherent point drift with joint scale, rotation, and translation optimization | Pose-matched geometric baseline |
| SSM-CPD | CPD deformation parameterized in statistical-shape-model coefficient space | Biology-guided robust estimator |
| SSM-CPD + refinement | SSM-CPD followed by unconstrained CPD refinement | Test whether local refinement improves fit and morphometric covariance |
| Topt + SSM-CPD + refinement | Template optimization followed by SSM-CPD and CPD refinement | Diagnostic severe-perturbation condition (“template-optimized” in Fig. 3B) |
| Integrated recovery | Template optimization, rigid pose recovery, SSM-CPD, and CPD refinement | Tail-rescue estimator for severe initialization failures |

CPD, coherent point drift; SSM, statistical shape model; Topt, template optimization.

To distinguish pose optimization from registration formulation, a focused 2 × 2 attribution analysis compared conventional CPD and the complete SSM-CPD estimator with pose either fixed or jointly optimized. All four conditions were evaluated for every held-out specimen (skulls and molars) or individual (vertebrae) under all three initialization regimes. The SSM-CPD formulation was treated as one integrated estimator comprising coefficient-space parameterization and covariance-weighted regularization.

Parameters were fixed before evaluation and were not tuned per specimen. All methods used *β* = 2, outlier weight *w* = 0.1, a maximum of 250 iterations, and tolerance 10^−6^. Conventional CPD and the refinement stage additionally used motion-coherence weight *α* = 2 and a low-rank approximation of the CPD kernel with at most 80 eigenvectors. SSM-CPD used the 95% variance subspace described above and regularization strength *λ*_reg_ = 0.01. Responsibility updates used a sparse k-d tree approximation with 10 nearest neighbors (Supplementary Note S1). All three primary estimators jointly optimized scale, rotation, and translation and used the same point-density parameter (1.3), which controls the density of the subsampled target point cloud.

Before registration, source and target coordinates were expressed in a shared target-scaled coordinate system in which the target bounding-box diagonal equaled 20, following Porto *et al*. (2021). Registration outputs were mapped back to the target coordinate system before landmark evaluation. Within SSM-CPD, target points were additionally centered and divided by their root-mean-square radius. The source configuration, mean shape, and SSM basis were transformed consistently and denormalized after convergence. This numerical normalization does not alter the reported percentage-of-bounding-box errors.

In SSM-CPD (Supplementary Note S1), the CPD E-step computes posterior responsibilities between target points and transformed atlas points. The M-step then solves for the SSM coefficients by minimizing a weighted squared-error term together with an eigenvalue-weighted coefficient penalty. In compact form, the coefficient update solves the regularized normal equation

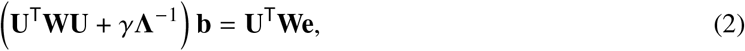

where **W** contains the posterior responsibility weights, **e** is the residual in the inverse-similarity frame, and *γ* = *λ*_reg_*σ*^2^/*s*^2^ combines the regularization strength *λ*_reg_, the current CPD variance *σ*^2^, and the similarity scale *s*.

### 2.6 Integrated recovery estimator for severe perturbation

We developed and evaluated an integrated recovery estimator for targets subjected to severe perturbation (Table 2; Supplementary Note S2). Standard atlas-based landmark transfer treats the source template as fixed, but a fixed template may lie too far from a perturbed or morphologically divergent target for registration to recover reliably. The estimator searches the learned SSM space for a target-compatible, biologically plausible template, uses that template for feature-based global pose recovery, and then applies SSM-CPD + refinement. Although designed as a rescue procedure, it was applied to every severely perturbed target rather than only to cases that failed, so that outcome-dependent selection could not inflate its apparent benefit.

Rigid pose recovery within the integrated estimator used feature-based alignment: point clouds were downsampled, normals and FPFH descriptors were computed, candidate rigid transformations were estimated by RANSAC, and local ICP refinement was applied (Fischler & Bolles, 1981; Rusu *et al*., 2009; Besl & McKay, 1992; Rusinkiewicz & Levoy, 2001). Its purpose is to place the optimized biological template and target within a recoverable pose basin before SSM-CPD and CPD refinement.

### 2.7 Landmark transfer accuracy

Registration was performed on the dense atlas correspondences. After the dense atlas points were warped to the target, sparse atlas landmarks were transferred by thin-plate-spline interpolation from the original to the warped dense atlas, with smoothing parameter *λ*_TPS_ = 0 (Bookstein, 1989). This yielded one predicted sparse landmark configuration for each specimen, method, and initialization regime.

Predicted sparse landmarks were compared with expert landmarks using per-landmark Euclidean error, summarized for each specimen as mean, median, RMSE, and maximum error, together with similarity-aligned Procrustes RMSE, runtime, and convergence metadata. The primary accuracy measure is each specimen’s median landmark error expressed as a percentage of the bounding-box diagonal of the subsampled target point cloud used for registration, which normalizes accuracy across anatomical structures of different sizes. Tail diagnostics report the median, 90th percentile (P90), 95th percentile (P95), and maximum of this measure across specimens, together with failure rates above 6% of the target bounding-box diagonal.

### 2.8 Shape preservation and covariance analysis

Pointwise landmark error is necessary, but not sufficient, for evaluating automated morphometric correspondence. A method can achieve low median error while distorting the multivariate shape structure used in downstream analyses (Fruciano, 2016; Grabowski & Porto, 2017; Percival *et al*., 2019; Savriama & Tautz, 2022). We therefore also evaluated the aligned regime with shape-preservation diagnostics. Manual and automated sparse landmark configurations were aligned by generalized Procrustes analysis (GPA), and automated configurations were compared with manual configurations in manual GPA/PCA space (Rohlf & Slice, 1990; Dryden & Mardia, 1998; Mitteroecker & Gunz, 2009).

The shape analysis quantified preservation of pairwise Procrustes distances, manual PC1 and PC2 scores, PC-score correlations for leading manual PCs, and covariance structure in manual PC space. For covariance diagnostics, both manual and automated configurations were projected into manual GPA/PCA space, enough manual PCs were retained to explain 80% of manual shape variance (PC80), and automated covariance was compared with manual covariance using covariance-trace ratios and PC-score correlations. This PC-space diagnostic is used because full coordinate covariance can be unstable when manual shape variance is small relative to coordinate dimensionality.

### 2.9 Statistical analysis and uncertainty

Registration error is reported as a percentage of the target bounding-box diagonal. Primary tail summaries use medians, P90/P95 quantiles, and prespecified failure thresholds. For matched method contrasts, positive reductions indicate lower error for the second method, and the win rate is the percentage of independent units for which the second method had lower error. Percentile bootstrap confidence intervals use 2,000 deterministic resamples, with specimens as independent units for skulls and molars and 85 biological-individual medians for vertebrae; paired Wilcoxon signed-rank tests use the same units. Shape-preservation intervals likewise use 2,000 biological-unit bootstrap resamples, with vertebral resampling clustered by individual. Because pairwise Procrustes distances are non-independent, their correlations are treated as descriptive association measures and ordinary correlation *P*-values are not reported.

### 2.10 Software implementation and availability

The SSM-CPD estimator is implemented in the biocpd library (https://github.com/agporto/biocpd). The primary registration estimators and the integrated recovery workflow are exposed through the MorphoWeave 3D Slicer extension (https://github.com/agporto/SlicerMorphoWeave). MorphoWeave supports atlas construction, statistical-shape-model building, and single- or batch-mode landmark prediction, allowing the benchmarked methods to be applied to new datasets without custom workflow code.

## 3 Results

### 3.1 SSM regularization reduces initialization failure

The primary robustness benchmark includes all held-out specimens across three datasets, three initialization regimes, and three primary methods. The experimental design is summarized in Fig. 1, and the primary robustness results are shown in Fig. 2 and Table 3. Under severe perturbation, CPD + joint similarity retained high median errors: 17.57% of bounding-box diagonal for skulls, 12.73% for molars, and 20.88% for vertebrae, and 100.0%, 83.5%, and 100.0% of specimens, respectively, exceeded the 6% failure threshold.

**Figure 2:**
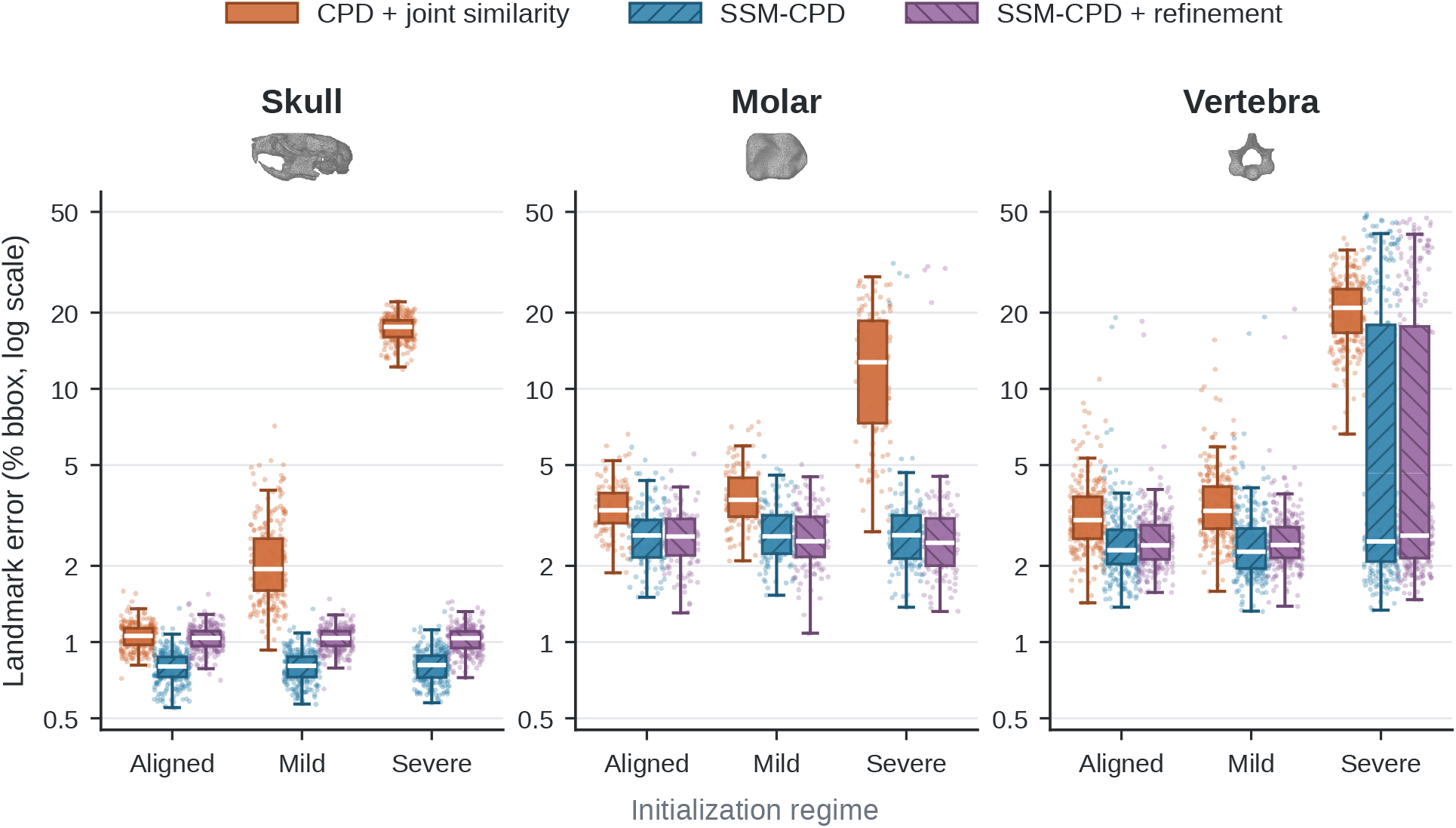
Registration error under aligned, mild, and severe initialization regimes. All three estimators jointly optimize a similarity transformation, so differences compare deformation models rather than unequal pose handling. Boxes show medians and interquartile ranges; whiskers extend to 1.5× the interquartile range; points are specimens. Vertebral points comprise 254 observations from 85 biological individuals, and inferential summaries use individual-level medians. The complete 2 × 2 pose-attribution analysis is provided in Supplementary Fig. S1.

**Table 3:** Severe-perturbation tail-risk and runtime summaries at the descriptive specimen level.

| Dataset | Method | *n* | Median | P90 | P95 | Failure > 6% | Time (s) |
| --- | --- | --- | --- | --- | --- | --- | --- |
| Skull | CPD + joint similarity | 249 | 17.57 | 19.75 | 20.38 | 100.0% | 4.19 |
| Skull | SSM-CPD | 249 | 0.81 | 0.95 | 0.99 | 0.0% | 1.82 |
| Skull | SSM-CPD + refinement | 249 | 1.04 | 1.16 | 1.23 | 0.0% | 4.08 |
| Skull | Topt + SSM-CPD + refinement | 249 | 1.03 | 1.20 | 1.26 | 0.0% | 14.12 |
| Skull | Integrated recovery | 249 | 1.04 | 1.20 | 1.26 | 0.0% | 14.92 |
| Molar | CPD + joint similarity | 115 | 12.73 | 22.73 | 25.50 | 83.5% | 2.68 |
| Molar | SSM-CPD | 115 | 2.64 | 3.85 | 4.93 | 3.5% | 1.30 |
| Molar | SSM-CPD + refinement | 115 | 2.47 | 3.76 | 4.03 | 3.5% | 3.60 |
| Molar | Topt + SSM-CPD + refinement | 115 | 2.57 | 3.76 | 4.37 | 3.5% | 22.32 |
| Molar | Integrated recovery | 115 | 2.42 | 3.64 | 4.18 | 2.6% | 23.59 |
| Vertebra | CPD + joint similarity | 254 | 20.88 | 28.17 | 31.04 | 100.0% | 5.79 |
| Vertebra | SSM-CPD | 254 | 2.50 | 37.46 | 41.35 | 28.0% | 2.37 |
| Vertebra | SSM-CPD + refinement | 254 | 2.63 | 35.99 | 40.77 | 27.6% | 6.97 |
| Vertebra | Topt + SSM-CPD + refinement | 254 | 2.61 | 35.49 | 38.74 | 28.0% | 45.52 |
| Vertebra | Integrated recovery | 254 | 2.44 | 3.62 | 4.10 | 4.3% | 50.08 |

Values summarize, across specimens, each specimen’s median sparse-landmark error as a percentage of the bounding-box diagonal of the subsampled target point cloud. Failure rate is the descriptive fraction of specimens with error > 6%. Time is the median wall-clock time of the complete pipeline per specimen. Topt, template optimization. Vertebral inferential comparisons use 85 biological-individual medians.

SSM-CPD largely eliminated this failure mode in skulls and molars and substantially reduced it in vertebrae. Under severe perturbation, median error decreased to 0.81% in skulls, 2.64% in molars, and 2.50% in vertebrae. SSM-CPD + refinement produced median errors of 1.04%, 2.47%, and 2.63%. Relative to CPD + joint similarity, SSM-CPD + refinement reduced paired severe-perturbation error by medians of 16.51 percentage points in skulls, 10.02 in molars, and 15.79 in vertebrae, with win rates of 100.0%, 97.4%, and 84.7%, respectively. Vertebral paired inference used 85 individual-level medians; improvement was strongly supported in every dataset (largest *P* = 1.68 × 10^−13^; Table 4).

**Table 4:** Matched biological-unit paired effects under severe perturbation.

| Dataset | Comparison | *n* | Median $\Delta$ | P90 $\Delta$ | Win rate | Wilcoxon *P* |
| --- | --- | --- | --- | --- | --- | --- |
| Skull | CPD + joint similarity → SSM-CPD + refinement | 249 | 16.51 | 18.72 | 100.0% | $1.35 \times 10^{-42}$ |
| Skull | SSM-CPD + refinement → integrated recovery | 249 | -0.03 | 0.06 | 37.3% | $1.47 \times 10^{-5}$ |
| Molar | CPD + joint similarity → SSM-CPD + refinement | 115 | 10.02 | 19.31 | 97.4% | $1.02 \times 10^{-19}$ |
| Molar | SSM-CPD + refinement → integrated recovery | 115 | -0.02 | 0.60 | 47.8% | 0.633 |
| Vertebra | CPD + joint similarity → SSM-CPD + refinement | 85 | 15.79 | 21.40 | 84.7% | $1.68 \times 10^{-13}$ |
| Vertebra | SSM-CPD + refinement → integrated recovery | 85 | 0.09 | 24.56 | 68.2% | $3.24 \times 10^{-5}$ |

Reductions are baseline minus comparison-method error in percentage points; positive values favor the second method. Win rate is the percentage of units for which the second method had lower error. P90 is the 90th percentile of paired reductions, not the comparison method's P90 error. Skull and molar units are specimens; vertebral inference uses 85 biological-individual medians.

To separate similarity optimization from the complete SSM formulation, a 2 × 2 attribution analysis crossed deformation model (unconstrained CPD versus SSM-CPD) with pose treatment (fixed versus jointly optimized). Under severe perturbation and matched joint similarity, the SSM formulation reduced median error by 16.72 percentage points in skulls (95% biological-unit bootstrap CI 16.34–16.91), 9.52 percentage points in molars (95% CI 7.65–11.13), and 16.09 percentage points in vertebrae (95% CI 15.30–17.43). Joint similarity optimization also reduced the error of conventional CPD, by 1.11, 7.65, and 0.70 percentage points, respectively, but this pose effect was smaller than the formulation effect in every dataset. The full 2 × 2 results and descriptive summaries are reported in Supplementary Fig. S1 and Supplementary Tables S1–S2.

### 3.2 Tail-risk diagnostics reveal failures that medians obscure

Tail-risk curves plot the fraction of held-out specimens whose sparse landmark error exceeds each threshold, making high-error failures visible even when median error is low (Fig. 3A; Table 3). For CPD + joint similarity, severe-perturbation P90 errors were 19.75% in skulls, 22.73% in molars, and 28.17% in vertebrae. Both SSM-CPD variants reduced this tail sharply in skulls and molars. For SSM-CPD + refinement, the corresponding P90 errors were 1.16% and 3.76%, and 0.0% and 3.5% of specimens, respectively, exceeded the 6% failure threshold.

**Figure 3:**
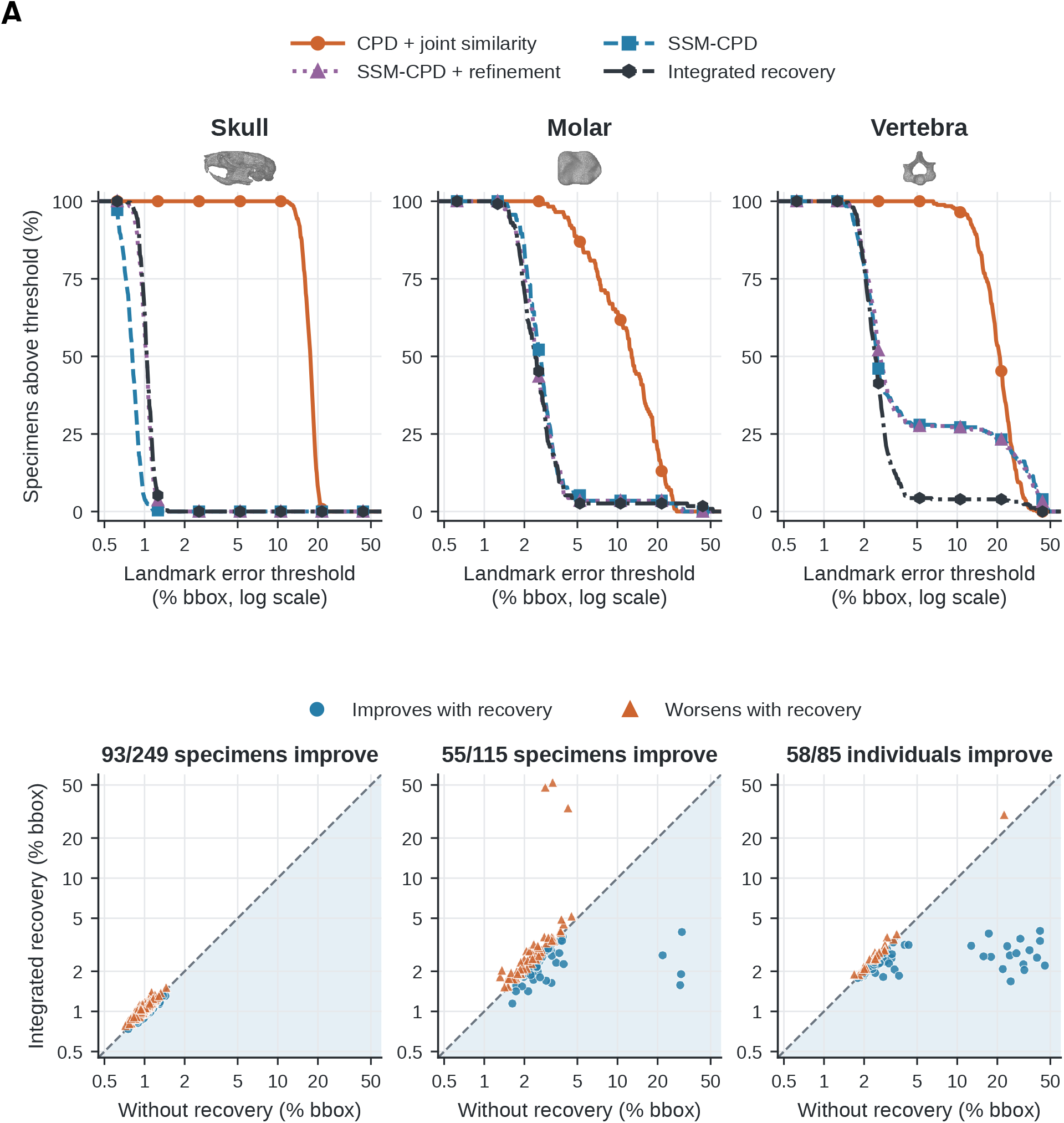
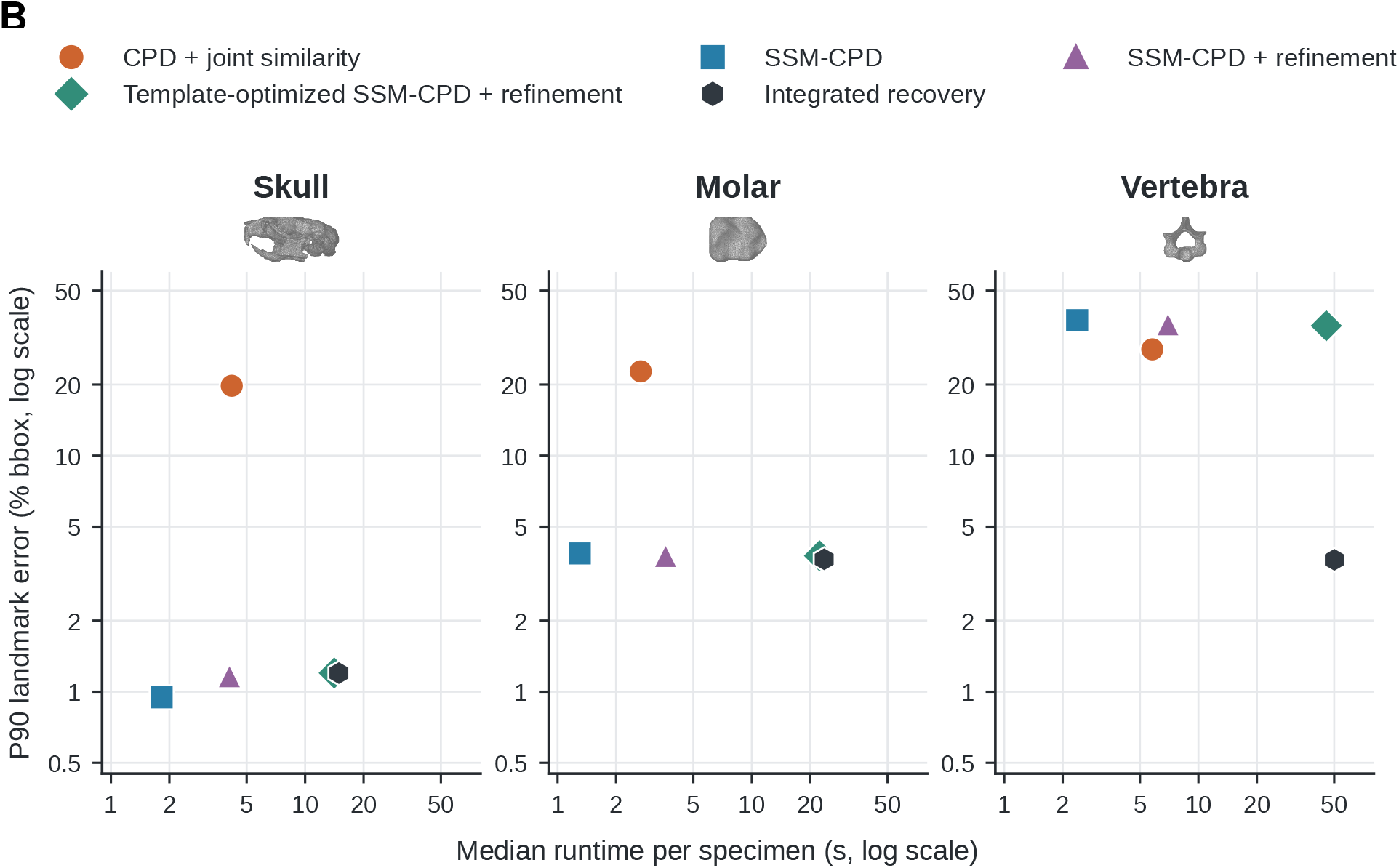
Tail-risk behavior, matched response, and computational trade-off under severe perturbation. **(A)** Top: percentage of specimens whose error exceeds each threshold for CPD + joint similarity, SSM-CPD, SSM-CPD + refinement, and integrated recovery. Bottom: paired responses to integrated recovery at the independent-unit level (specimens for skulls and molars, individual-level medians for vertebrae); points in the shaded region below the diagonal improve. Axes are identical across datasets. **(B)** Median runtime per specimen versus P90 landmark error for the primary estimators, template-optimized SSM-CPD + refinement (Topt + SSM-CPD + refinement in Table 2), and integrated recovery.

Vertebrae were more difficult. SSM-CPD and SSM-CPD + refinement had low median errors, but their P90 errors remained 37.46% and 35.99%, and 28.0% and 27.6% of vertebrae, respectively, exceeded the 6% threshold. These cases motivate the integrated recovery analysis in the next section (Fig. 3A, bottom row). Fig. 3B relates tail behavior to computational cost: SSM-CPD was the fastest estimator in every dataset (median 1.30–2.37 s per specimen), whereas integrated recovery required 14.92–50.08 s (Table 3).

### 3.3 Integrated recovery rescues the vertebral high-error tail

The severe-perturbation recovery comparison evaluated SSM-CPD + refinement against integrated recovery for all 618 held-out targets across the three datasets (Fig. 4; Table 4).

**Figure 4:**
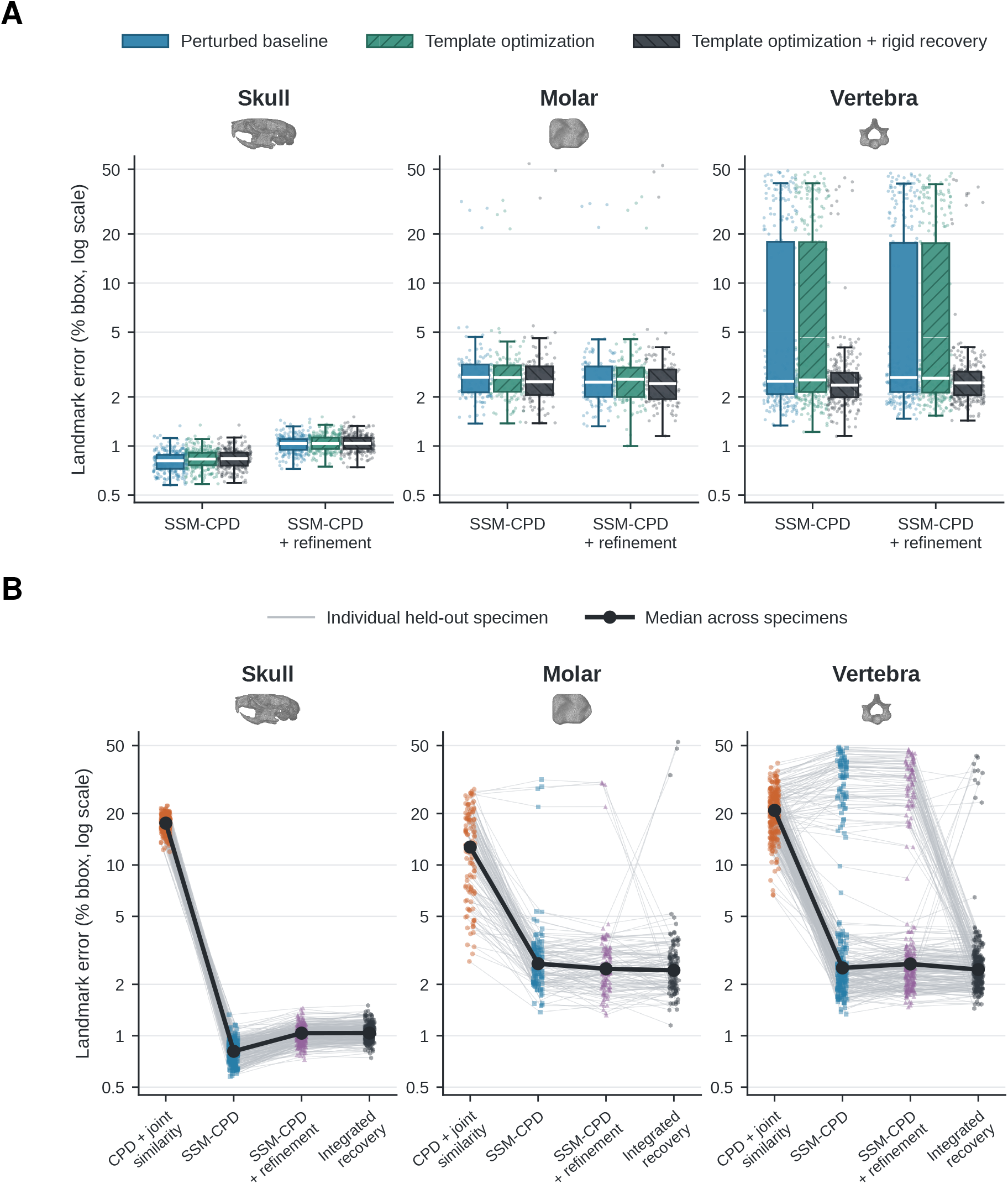
Integrated recovery under severe perturbation. **(A)** Specimen-level error distributions for SSM-CPD and SSM-CPD + refinement at the perturbed baseline, after template optimization, and after template optimization with rigid recovery; the dark box in the SSM-CPD + refinement group is integrated recovery. **(B)** Matched specimen-level trajectories across CPD + joint similarity, SSM-CPD, SSM-CPD + refinement, and integrated recovery; thin lines are held-out specimens and the thick line joins condition-wise medians. Biological-unit inference is reported separately in Table 4, using specimens for skulls and molars and 85 individual-level medians for vertebrae.

In vertebrae, integrated recovery sharply reduced the high-error tail. The specimen-level P90 error decreased from 35.99% to 3.62%, and the proportion of vertebrae above the 6% threshold decreased from 27.56% to 4.33%. At the level of biological individuals, 58/85 individuals improved, the median paired reduction was 0.09 percentage points, and the 90th percentile of paired reductions was 24.56 percentage points. Residual risk remained: 11 of 254 vertebrae (4.33%) still exceeded the 6% threshold after recovery, with a maximum error of 43.30%. Recovery rescued 65 of the 70 vertebral failures under SSM-CPD + refinement, but six vertebrae that had not failed without recovery exceeded the threshold after it.

Matched responses show why integrated recovery should be interpreted as conditional rather than universally beneficial (Figs. 3A and 4B; Table 4). Relative to SSM-CPD + refinement, 93/249 skull specimens, 55/115 molar specimens, and 58/85 vertebral individuals improved. The median paired change was negligible in skulls (−0.03 percentage points; statistically detectable but of no practical consequence) and in molars (−0.02; *P* = 0.63), whereas the benefit was concentrated in the vertebral tail. In molars, recovery rescued all four failures under SSM-CPD + refinement but introduced three new failures (Supplementary Table S4). The largest rescue among vertebrae reduced error from 47.35% to 2.22%, and eight additional vertebrae improved by more than 40 percentage points. Because recovery also increased median runtime roughly four-to sevenfold (Table 3), it is most useful for difficult anatomical systems or for cases flagged by failure diagnostics.

### 3.4 Automated landmarks preserve major morphometric structure

GPA and PCA were used to evaluate whether automated landmarks preserve biological shape structure in the aligned regime (Fig. 5; Table 5). Pairwise Procrustes-distance preservation was moderate to high across datasets. In skulls, the correlation between manual and automated pairwise distances was 0.84 for CPD + joint similarity, 0.91 for SSM-CPD, and 0.88 for SSM-CPD + refinement. The corresponding values were 0.76, 0.86, and 0.90 in molars, and 0.80, 0.85, and 0.90 in vertebrae.

**Figure 5:**
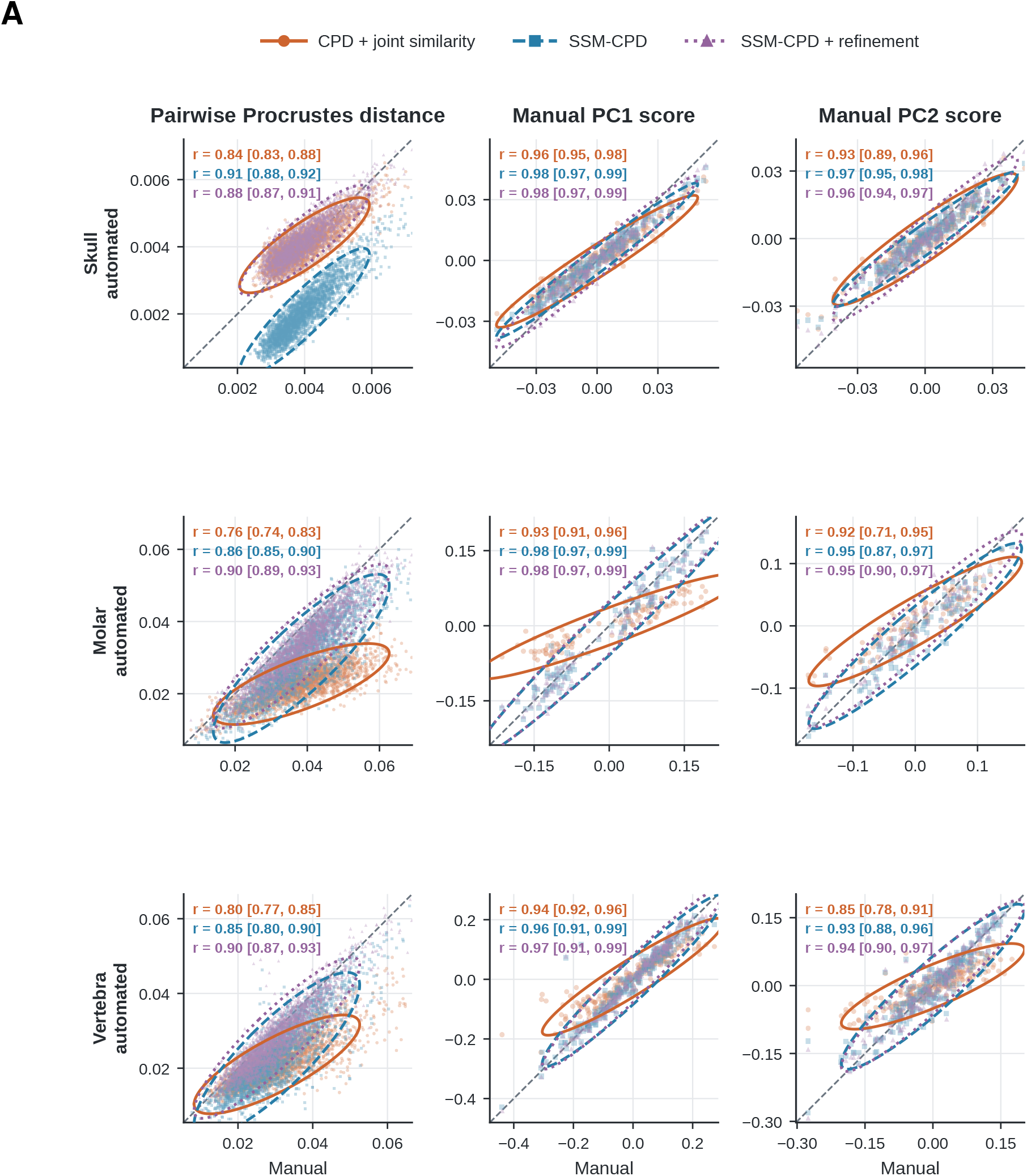
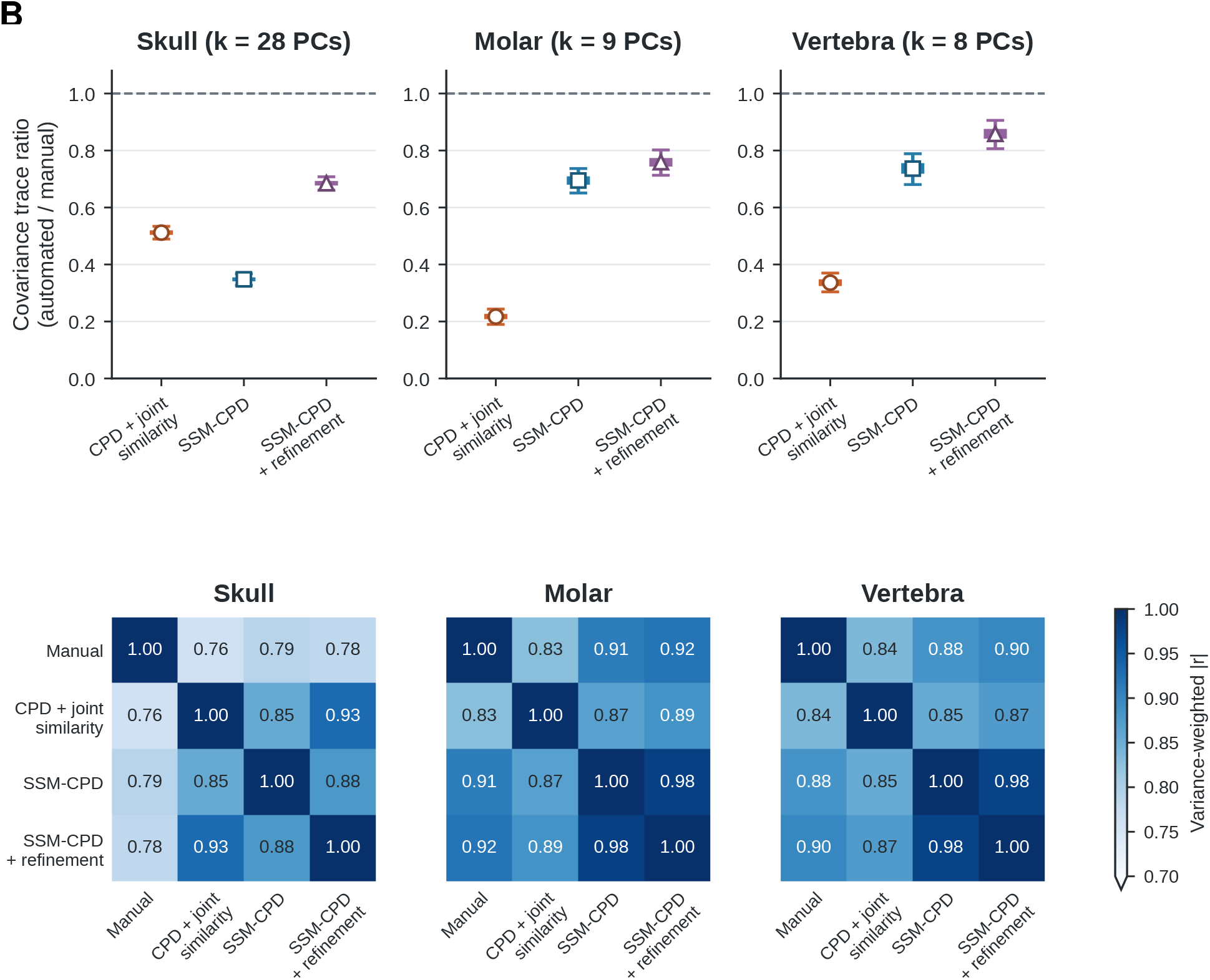
Preservation of morphometric structure in the aligned regime. **(A)** Manual configurations compared with predictions from CPD + joint similarity, SSM-CPD, and SSM-CPD with refinement using pairwise Procrustes distances and scores on the first two manual GPA/PCA axes. Ellipses enclose 95% of each method’s points under a bivariate normal approximation; values are correlations with 95% bootstrap intervals; dashed lines mark perfect preservation. **(B)** Top: bootstrap distributions of PC80 covariance-trace ratios (markers, bootstrap medians; thick bars, interquartile ranges; whiskers, 95% intervals; dashed line, ratio of 1). Bottom: variance-weighted absolute PC-score correlations among manual and automated configurations after projection into manual GPA/PCA space. Vertebral uncertainty is clustered by biological individual.

**Table 5:** Shape-preservation diagnostics in the aligned regime.

| Dataset | Method | *n* spec./units | Pairwise *r* | PC1 *r* | PC2 *r* | PC80 trace |
| --- | --- | --- | --- | --- | --- | --- |
| Skull | CPD + joint similarity | 249/249 | 0.84 | 0.96 | 0.93 | 0.56 |
| Skull | SSM-CPD | 249/249 | 0.91 | 0.98 | 0.97 | 0.35 |
| Skull | SSM-CPD + refinement | 249/249 | 0.88 | 0.98 | 0.96 | 0.74 |
| Molar | CPD + joint similarity | 115/115 | 0.76 | 0.93 | 0.92 | 0.23 |
| Molar | SSM-CPD | 115/115 | 0.86 | 0.98 | 0.95 | 0.71 |
| Molar | SSM-CPD + refinement | 115/115 | 0.90 | 0.98 | 0.95 | 0.78 |
| Vertebra | CPD + joint similarity | 254/85 | 0.80 | 0.94 | 0.85 | 0.34 |
| Vertebra | SSM-CPD | 254/85 | 0.85 | 0.96 | 0.93 | 0.75 |
| Vertebra | SSM-CPD + refinement | 254/85 | 0.90 | 0.97 | 0.94 | 0.87 |

Pairwise *r* is the Pearson correlation between manual and predicted pairwise Procrustes distances. PC correlations are component-wise correlations in manual GPA/PCA space. PC80 trace is the predicted/manual covariance-trace ratio. Vertebral inference uses 85 biological individuals.

The leading shape axes were strongly preserved. PC1 score correlations for CPD + joint similarity, SSM-CPD, and SSM-CPD + refinement were 0.96, 0.98, and 0.98 for skulls, 0.93, 0.98, and 0.98 for molars, and 0.94, 0.96, and 0.97 for vertebrae. PC2 correlations were also high, although lower than PC1 in several conditions (Table 5).

Covariance preservation revealed a trade-off not captured by pointwise error or PC-score correlations. In skulls, SSM-CPD compressed covariance in manual PC space: its PC80 covariance-trace ratio (0.35) was lower than that of CPD + joint similarity (0.56), and refinement raised it to 0.74. In molars and vertebrae, by contrast, SSM-CPD improved covariance preservation relative to CPD + joint similarity (from 0.23 to 0.71 and from 0.34 to 0.75, respectively), and refinement increased the ratio further, to 0.78 and 0.87. Aligned factorial controls produced similar shape-preservation estimates whether pose was fixed or jointly optimized, indicating that these morphometric differences reflect the deformation model rather than pose handling (Supplementary Fig. S2; Supplementary Table S3).

## 4 Discussion

Anatomical correspondence defines the variables of geometric morphometrics (Bookstein, 1991; Mitteroecker & Schaefer, 2022), and as three-dimensional imaging and collection digitization expand morphometric datasets, establishing it at scale has become a computational problem (Hedrick *et al*., 2020; Blackburn *et al*., 2024). Manual landmarking remains the reference standard but is difficult to scale and is itself subject to observer error (Fruciano, 2016), whereas automated landmark transfer depends on registration algorithms whose assumptions are geometric rather than biological. Our results show that automated 3D landmark transfer becomes more robust when registration is constrained not only by geometric fit but also by the empirical structure of biological shape variation.

This work integrates three related traditions. In organismal morphometrics, landmark-free surface comparison, automated pseudolandmarks, globally consistent correspondence, and data-driven landmark sampling have extended shape analysis beyond manually placed sparse landmarks (Boyer *et al*., 2011, 2015; Gao *et al*., 2018, 2019; Koehl & Hass, 2015). In computer vision and medical image analysis, statistical shape models and point distribution models have long constrained segmentation, registration, and reconstruction to plausible regions of learned shape space (Cootes *et al*., 1995; Heimann & Meinzer, 2009). In point-set registration, probabilistic methods such as coherent point drift treat correspondence as a latent-variable problem and estimate transformations while regularizing deformation smoothness (Myronenko & Song, 2010); related work has also coupled Gaussian-mixture correspondence with statistical shape priors (Hirose, 2017). The present study develops these ingredients into a complete, reusable framework for automated 3D biological landmark transfer. Homologous dense correspondences define the admissible deformation subspace, population covariance defines its precision structure, similarity pose is estimated jointly, and optional local refinement and integrated recovery extend practical performance. The contribution is therefore both algorithmic and operational: a coefficient-space SSM-CPD estimator, a target-adaptive recovery estimator, and open-source software that exposes both within an atlas-based morphometric workflow.

The biology-guided framework is important because the statistical shape model is not simply a post hoc dimensionality reduction. Instead, it changes the registration problem. In unconstrained CPD, deformation is guided by geometric proximity and motion coherence. In SSM-CPD, the deformation is expressed through coefficients of a population shape model and penalized according to the inverse eigenvalues of observed biological variation. The estimator therefore treats a deformation along a common biological mode differently from an equally large deformation along a rare or unsupported direction. In this sense, dense biological correspondence becomes an active prior on anatomical plausibility. This connects morphometric registration to a broader class of knowledge-guided machine-learning approaches, in which scientific structure is incorporated directly into the model or inference procedure rather than imposed only after prediction (Karpatne *et al*., 2017, 2024; Lürig *et al*., 2024).

Our empirical results support this view while clarifying the source and magnitude of the improvement. Under severe perturbation, SSM-CPD + refinement reduced paired error relative to CPD + joint similarity by medians of 16.51 percentage points in skulls, 10.02 in molars, and 15.79 in vertebrae. Joint similarity optimization improved conventional CPD, especially in molars, but the complete SSM formulation retained a large advantage when pose treatment was matched. The remaining advantage can therefore be attributed to the SSM formulation rather than to unequal pose handling.

The results also show why robustness should be evaluated with tail-aware diagnostics. Median landmark error is useful, but it can obscure the failure modes that matter most in large comparative datasets. In the vertebral dataset, for example, median error suggested that SSM-CPD performed well under severe perturbation, whereas tail-risk curves and high-percentile runtime–accuracy summaries revealed that more than a quarter of vertebrae still failed catastrophically. In large morphometric studies, even rare registration failures can bias estimates of disparity, covariance structure, allometry, group separation, or downstream genetic association, and they can impose substantial manual-cleaning costs (Fruciano, 2016; Percival *et al*., 2019; Savriama & Tautz, 2022). Automated landmarking methods should therefore be evaluated not only by central tendency, but also by the frequency, severity, and detectability of their failures. Practical image-analysis workflows can retain expert oversight by allowing users to inspect and filter automated segmentations or interactively correct predicted landmarks (Di Martino *et al*., 2023; Quintana *et al*., 2026).

The recovery results further clarify the role of initialization in biology-guided registration. Recovery is evaluated here as an integrated sequence: the learned SSM is searched for a target-compatible template, feature-based alignment recovers pose, and SSM-CPD + refinement estimates correspondence. In skulls and molars, where SSM-CPD already performed well, recovery added runtime with negligible typical change and, in molars, introduced a few new failures. In vertebrae, it rescued most high-error cases without eliminating every severe failure. This suggests a conditional workflow: use SSM-CPD as the default robust estimator and deploy integrated recovery when severe pose error, high morphological disparity, or diagnostic summaries indicate elevated failure risk.

The shape-preservation analyses add a second layer of insight. Landmarking accuracy and morphometric utility are related, but not identical. A method can place individual landmarks close to expert annotations while still altering the covariance structure used for downstream comparative analyses. Pure SSM regularization stabilizes correspondence, but it can also compress variation when the prior is strong relative to the target sample, as it did in skulls. CPD refinement restored more of the manual covariance structure, raising covariance-trace ratios in all three datasets. This trade-off argues against evaluating automated landmarking solely with pointwise error. Previous comparisons show that similar landmark errors can have different consequences for mean shape, covariance, and downstream biological signal (Percival *et al*., 2019; Savriama & Tautz, 2022). For morphometric applications, accuracy, tail risk, and preservation of shape-space structure should be reported together.

Several limitations should be noted. First, the SSM can only represent variation present in the training correspondences. If the training sample does not span the target distribution, the prior may bias registration toward known shapes and underrepresent novel morphology. Second, dense correspondences are treated as biologically meaningful inputs to the model; choices made during surface sampling and correspondence construction will propagate into the estimated shape prior (Bardua *et al*., 2019; Gao *et al*., 2018). Third, the perturbation regimes used here are controlled stress tests rather than a complete representation of real-world failure modes. They were designed to isolate initialization sensitivity, but future evaluations should include messier sources of error such as incomplete meshes, segmentation artifacts, damaged specimens, and taxonomic extrapolation beyond the training distribution. Fourth, because subspace restriction and covariance weighting are integrated in SSM-CPD, and the recovery stages were evaluated together, our experiments do not separate the contributions of these components. Finally, the comparisons isolate the effect of the SSM prior within CPD; we did not benchmark other automated landmarking approaches, such as multi-template or landmark-free methods, and the expert landmarks used as the reference standard carry their own observer error (Zhang *et al*., 2022; Boyer *et al*., 2011; Fruciano, 2016).

More broadly, these results argue that automated landmarking should be treated as an inferential component of morphometric analysis rather than as a neutral preprocessing step (Fruciano, 2016; Mitteroecker & Schaefer, 2022). Registration algorithms encode assumptions about which correspondences are plausible, which deformations are permitted and which errors are likely. The advantage of SSM-CPD is that these assumptions are made biological, explicit, and empirically testable. By embedding observed shape covariance into the correspondence estimator, biology-guided registration provides a route toward automated morphometrics that is not only scalable, but also better aligned with the anatomical structure that comparative biologists seek to measure. The open-source implementations in biocpd and MorphoWeave (3D Slicer) make the estimator and recovery workflow available as reusable components for morphometric analysis of large, complex biological datasets.

## 5 Conclusions

We developed, openly implemented, and benchmarked a biology-guided CPD framework for automated landmark transfer in which dense anatomical correspondence and population shape covariance are incorporated directly into the registration estimator. Across mouse skulls, primate molars, and squamate vertebrae, the complete SSM-CPD estimator reduced severe-initialization failures relative to conventional CPD under equivalent joint similarity optimization. A focused factorial analysis showed that this advantage was not explained by pose handling alone. The integrated recovery estimator was most valuable as a conditional strategy for the vertebral high-error tail. Shape-preservation diagnostics further showed complementary contributions of SSM regularization and CPD refinement: the prior stabilizes correspondence, while refinement can improve covariance preservation. Together, these results support biology-guided registration as a robust foundation for scalable 3D morphometrics. The open-source implementations in biocpd and MorphoWeave provide a practical path to adoption.

## Author contributions

Arthur Porto: Conceptualization, Methodology, Analysis, Writing – original draft, Supervision and Project administration. Katherine A. Wolcott: Data curation, Methodology, Analysis, Visualization and Writing – review and editing. Kelsey L. Fenner: Data curation and Writing – review and editing. Caleb Gordon: Data curation and Writing – review and editing. Breanna Shi: Methodology and Writing – review and editing. Edward L. Stanley: Resources, Data curation and Writing – review and editing. All authors contributed to the interpretation of the results, reviewed the manuscript, and approved the submitted version.

## Acknowledgements

We thank Murat Maga, Sara Rolfe, Sarah Du Plessis, Christy Hipsley, Natasha Vitek, and Eva Herbst for discussions and contributions to the initial testing of the MorphoWeave workflow. We thank Diogo Melo, Jason Wolf, and Gabriel Marroig for sharing the mouse models used in this study. We thank Osamu Hirose, Nikita Angarski, Omar Moursy, and Steve Foryoung for initial discussions on the coherent point drift derivation.

## Use of generative AI

OpenAI’s GPT-5.6 Pro and Anthropic’s Claude Fable 5 were used as coding assistants and to assist with language editing. All suggested text and code were reviewed by the authors, and the corresponding author takes responsibility for the final content.

