## Supplemental Material for "Biology-guided machine learning for robust anatomical correspondence across biodiversity datasets"

### Supplementary Note S1: Derivation of SSM-regularized coherent point drift

#### S1.1 Scope and notation

This note derives the statistical-shape-model-regularized coherent point drift (SSM-CPD) estimator developed for the `biocpd` library and integrated into the MorphoWeave atlas-registration workflow. The method combines the CPD likelihood with a low-dimensional statistical shape model and an eigenvalue-weighted coefficient prior (Myronenko & Song, 2010; Cootes *et al.*, 1995; Heimann & Meinzer, 2009; Hirose, 2017). The derivation concerns the dense atlas correspondences used during registration; sparse anatomical landmarks are transferred only after the dense deformation has been estimated.

Let the target point cloud be

$$\mathcal{X} = \{\mathbf{x}_n\}_{n=1}^{N}, \qquad \mathbf{x}_n \in \mathbb{R}^D, \tag{S1.1}$$

and let the source atlas correspondence configuration be

$$\mathcal{Y} = \{\mathbf{y}_m\}_{m=1}^{M}, \qquad \mathbf{y}_m \in \mathbb{R}^D. \tag{S1.2}$$

In this study, $D = 3$, $\mathcal{X}$ is a subsampled target mesh, and $\mathcal{Y}$ is the dense atlas-correspondence cloud. The same derivation applies whether $\mathcal{Y}$ is the supplied atlas configuration or an explicitly supplied mean configuration.

The SSM is represented by a basis matrix and eigenvalue matrix

$$\mathbf{U} \in \mathbb{R}^{MD\times K}, \qquad \mathbf{\Lambda} = \mathrm{diag}(\lambda_1, \ldots, \lambda_K), \qquad \lambda_k > 0. \tag{S1.3}$$

The implementation accepts $\mathbf{U}$ as either an $M \times D \times K$ array or a flattened $MD \times K$ matrix. Let

$$\mathbf{b} = (b_1, \ldots, b_K)^\top \in \mathbb{R}^K \tag{S1.4}$$

denote the SSM coefficients. The constrained source configuration is

$$\mathbf{Z}(\mathbf{b}) = \mathbf{Y} + \mathrm{unvec}(\mathbf{U}\mathbf{b}), \tag{S1.5}$$

or, for source point $m$,

$$\mathbf{z}_m(\mathbf{b}) = \mathbf{y}_m + \mathbf{U}_m\mathbf{b}, \tag{S1.6}$$

where $\mathbf{U}_m \in \mathbb{R}^{D\times K}$ is the basis block associated with point $m$. The transformed source point is

$$\mathbf{T}_m(\mathbf{b}, \mathbf{R}, s, \mathbf{t}) = s\mathbf{R}\mathbf{z}_m(\mathbf{b}) + \mathbf{t}, \tag{S1.7}$$

where $\mathbf{R} \in \mathrm{SO}(D)$, $s > 0$ is an optional uniform scale, and $\mathbf{t} \in \mathbb{R}^D$. The implementation evaluates this expression as `s * (Z @ R.T) + t`; this is the row-vector convention corresponding to the column-vector notation above.

#### S1.2 Standard CPD likelihood

CPD treats the transformed source points as centroids of a Gaussian mixture (Myronenko & Song, 2010). Conditional on component $m$, target point $\mathbf{x}_n$ is generated as

$$p(\mathbf{x}_n \mid m) = (2\pi\sigma^2)^{-D/2} \exp\left[-\frac{\|\mathbf{x}_n - \mathbf{T}_m\|^2}{2\sigma^2}\right]. \tag{S1.8}$$

The mixture includes a uniform-noise component with weight $w \in [0, 1)$. The posterior responsibility that source centroid $m$ explains target point $n$ is

$$P_{mn} = \frac{\exp\left[-\|\mathbf{x}_n - \mathbf{T}_m\|^2/(2\sigma^2)\right]}{\sum_{j=1}^{M} \exp\left[-\|\mathbf{x}_n - \mathbf{T}_j\|^2/(2\sigma^2)\right] + c}, \tag{S1.9}$$

with

$$c = (2\pi\sigma^2)^{D/2} \frac{w}{1-w} \frac{M}{N}. \tag{S1.10}$$

The expectation step produces the sufficient statistics

$$P1_m = \sum_{n=1}^{N} P_{mn}, \tag{S1.11}$$

$$P^{\mathsf{T}}1_n = \sum_{m=1}^{M} P_{mn}, \tag{S1.12}$$

$$N_p = \sum_{m=1}^{M} \sum_{n=1}^{N} P_{mn}, \tag{S1.13}$$

$$PX_m = \sum_{n=1}^{N} P_{mn} \mathbf{x}_n. \tag{S1.14}$$

For every source point with non-zero responsibility mass, define its posterior-weighted target mean

$$\bar{\mathbf{x}}_m = \frac{PX_m}{P1_m}. \tag{S1.15}$$

The M-step can then be written in terms of these $M$ weighted target means rather than the full $M \times N$ correspondence matrix.

#### S1.3 Complete-data objective

Ignoring constants independent of the transformation, the CPD M-step minimizes

$$Q_{\text{data}}(\mathbf{b}, \mathbf{R}, s, \mathbf{t}) = \frac{1}{2\sigma^2} \sum_{m=1}^{M} \sum_{n=1}^{N} P_{mn} \|\mathbf{x}_n - s\mathbf{R}\mathbf{z}_m(\mathbf{b}) - \mathbf{t}\|^2. \tag{S1.16}$$

Using the posterior means, the parameter-dependent part is equivalently

$$Q_{\text{data}}(\mathbf{b}, \mathbf{R}, s, \mathbf{t}) = \frac{1}{2\sigma^2} \sum_{m=1}^{M} P1_m \left\| \bar{\mathbf{x}}_m - s\mathbf{R}\mathbf{z}_m(\mathbf{b}) - \mathbf{t} \right\|^2 . \tag{S1.17}$$

SSM-CPD adds the eigenvalue-weighted coefficient penalty

$$Q_{\text{prior}}(\mathbf{b}) = \frac{\lambda_{\text{reg}}}{2} \mathbf{b}^{\mathsf{T}} \mathbf{\Lambda}^{-1} \mathbf{b}. \tag{S1.18}$$

This is a Gaussian shape prior: coefficients along high-variance SSM modes are penalized less than coefficients along low-variance modes. The MAP objective is

$$Q = Q_{\text{data}} + Q_{\text{prior}}. \tag{S1.19}$$

Registration alternates among responsibility, SSM-coefficient, and optional similarity-parameter updates.

#### S1.4 Coefficient update for fixed similarity

Hold $(\mathbf{R}, s, \mathbf{t})$ fixed. The inverse similarity map applied to the posterior target mean is

$$\mathbf{i}_m = T^{-1}(\bar{\mathbf{x}}_m) = \frac{1}{s} \mathbf{R}^{\mathsf{T}} (\bar{\mathbf{x}}_m - \mathbf{t}). \tag{S1.20}$$

Under the row-vector convention this is $(\bar{\mathbf{x}}_m - \mathbf{t})\mathbf{R}/s$. Because rotations preserve Euclidean distances,

$$\left\| \bar{\mathbf{x}}_m - s\mathbf{R}\mathbf{z}_m(\mathbf{b}) - \mathbf{t} \right\|^2 = s^2 \left\| \mathbf{i}_m - \mathbf{z}_m(\mathbf{b}) \right\|^2 . \tag{S1.21}$$

Substitution into the MAP objective gives

$$Q(\mathbf{b}) = \frac{s^2}{2\sigma^2} \sum_{m=1}^{M} P1_m \left\| \mathbf{i}_m - \mathbf{y}_m - \mathbf{U}_m \mathbf{b} \right\|^2 + \frac{\lambda_{\text{reg}}}{2} \mathbf{b}^{\mathsf{T}} \mathbf{\Lambda}^{-1} \mathbf{b}. \tag{S1.22}$$

Multiplication by $2\sigma^2/s^2$ does not change the minimizer, so the coefficient update minimizes

$$J(\mathbf{b}) = \sum_{m=1}^{M} P1_m \left\| \mathbf{i}_m - \mathbf{y}_m - \mathbf{U}_m \mathbf{b} \right\|^2 + \gamma \mathbf{b}^{\mathsf{T}} \mathbf{\Lambda}^{-1} \mathbf{b}, \qquad \gamma = \frac{\lambda_{\text{reg}} \sigma^2}{s^2}. \tag{S1.23}$$

When scale is disabled, $s = 1$.

Let

$$\mathbf{e} = \operatorname{vec}(\mathbf{i}_1 - \mathbf{y}_1, \ldots, \mathbf{i}_M - \mathbf{y}_M) \in \mathbb{R}^{MD}, \tag{S1.24}$$

and define

$$\mathbf{W} = \operatorname{diag}(P1_1 \mathbf{I}_D, \ldots, P1_M \mathbf{I}_D) \in \mathbb{R}^{MD \times MD}. \tag{S1.25}$$

Then

$$J(\mathbf{b}) = (\mathbf{e} - \mathbf{U}\mathbf{b})^{\mathsf{T}} \mathbf{W} (\mathbf{e} - \mathbf{U}\mathbf{b}) + \gamma \mathbf{b}^{\mathsf{T}} \mathbf{\Lambda}^{-1} \mathbf{b}. \tag{S1.26}$$

Expanding and differentiating with respect to $\mathbf{b}$ yields

$$-2\mathbf{U}^{\mathsf{T}}\mathbf{W}\mathbf{e} + 2\mathbf{U}^{\mathsf{T}}\mathbf{W}\mathbf{U}\mathbf{b} + 2\gamma\mathbf{\Lambda}^{-1}\mathbf{b} = 0. \tag{S1.27}$$

Therefore,

$$\boxed{\left(\mathbf{U}^{\mathsf{T}}\mathbf{W}\mathbf{U} + \gamma\mathbf{\Lambda}^{-1}\right)\mathbf{b} = \mathbf{U}^{\mathsf{T}}\mathbf{W}\mathbf{e}} \tag{S1.28}$$

and

$$\hat{\mathbf{b}} = \left(\mathbf{U}^{\mathsf{T}}\mathbf{W}\mathbf{U} + \frac{\lambda_{\text{reg}}\sigma^2}{s^2}\mathbf{\Lambda}^{-1}\right)^{-1}\mathbf{U}^{\mathsf{T}}\mathbf{W}\mathbf{e}. \tag{S1.29}$$

This is the central computational difference from unconstrained deformable CPD: SSM-CPD estimates a low-dimensional coefficient vector regularized by a biological shape-covariance model.

#### S1.5 Direct correspondence to the implementation

The implementation computes

$$\bar{\mathbf{x}}_m = PX_m/P1_m, \tag{S1.30}$$

applies the inverse similarity transform, constructs $\mathbf{e} = \text{vec}(\mathbf{i}_m - \mathbf{y}_m)$, and forms

$$\mathbf{A} = \mathbf{U}^{\mathsf{T}}\mathbf{W}\mathbf{U} + \gamma\mathbf{\Lambda}^{-1}, \tag{S1.31}$$

$$\mathbf{r} = \mathbf{U}^{\mathsf{T}}\mathbf{W}\mathbf{e}. \tag{S1.32}$$

It solves $\mathbf{A}\mathbf{b} = \mathbf{r}$ by Cholesky factorization. The resulting deformation and updated source are

$$\mathbf{\Delta} = \text{unvec}(\mathbf{U}\hat{\mathbf{b}}), \qquad \mathbf{Y}_b = \mathbf{Y} + \mathbf{\Delta}. \tag{S1.33}$$

When similarity optimization is enabled, weighted Procrustes estimation (Rohlf & Slice, 1990) updates $(\mathbf{R}, s, \mathbf{t})$ and sets

$$T\mathbf{Y} = s\mathbf{R}\mathbf{Y}_b + \mathbf{t}. \tag{S1.34}$$

Otherwise, $T\mathbf{Y} = \mathbf{Y}_b$.

#### S1.6 Weighted Procrustes update for similarity parameters

Given $\mathbf{Y}_b = \{\mathbf{y}_m^b\}_{m=1}^M$, the implementation minimizes

$$\sum_{m=1}^{M} P1_m \left\|\bar{\mathbf{x}}_m - s\mathbf{R}\mathbf{y}_m^b - \mathbf{t}\right\|^2. \tag{S1.35}$$

Define weighted means

$$\boldsymbol{\mu}_x = \frac{1}{N_p}\sum_{m=1}^{M} P1_m\bar{\mathbf{x}}_m, \qquad \boldsymbol{\mu}_y = \frac{1}{N_p}\sum_{m=1}^{M} P1_m\mathbf{y}_m^b, \tag{S1.36}$$

and centered points

$$\mathbf{X}_m^c = \bar{\mathbf{x}}_m - \boldsymbol{\mu}_x, \qquad \mathbf{Y}_m^c = \mathbf{y}_m^b - \boldsymbol{\mu}_y. \tag{S1.37}$$

The weighted cross-covariance is

$$\mathbf{C} = \sum_{m=1}^{M} P1_m (\mathbf{Y}_m^c)^\top \mathbf{X}_m^c. \tag{S1.38}$$

Let

$$\mathbf{C} = \mathbf{U}_c \mathbf{\Sigma} \mathbf{V}_c^\top \tag{S1.39}$$

be its singular value decomposition, and define

$$\mathbf{S}_c = \operatorname{diag}\Big(1, \ldots, 1, \operatorname{sign}\det(\mathbf{U}_c \mathbf{V}_c^\top)\Big) \tag{S1.40}$$

to prevent reflection. Under the row-vector convention,

$$\mathbf{R} = \Big(\mathbf{U}_c \mathbf{S}_c \mathbf{V}_c^\top\Big)^\top. \tag{S1.41}$$

When scale is estimated,

$$s = \frac{\operatorname{tr}\big[(\mathbf{U}_c \mathbf{S}_c \mathbf{V}_c^\top)^\top \mathbf{C}\big]}{\sum_{m=1}^{M} P1_m \|\mathbf{Y}_m^c\|^2}. \tag{S1.42}$$

If scale is disabled, $s = 1$. Translation is

$$\mathbf{t} = \boldsymbol{\mu}_x - s\boldsymbol{\mu}_y \mathbf{R}^\top. \tag{S1.43}$$

These expressions correspond directly to the implementation's `_weighted_similarity_update` routine.

#### S1.7 Variance update

After updating the transformed source points, residual variance is updated by

$$\sigma^2 = \frac{1}{N_p D} \sum_{m=1}^{M} \sum_{n=1}^{N} P_{mn} \|\mathbf{x}_n - T\mathbf{Y}_m\|^2. \tag{S1.44}$$

The implementation evaluates the expanded form

$$\sigma^2 = \frac{xPx - 2\operatorname{tr}(PXY) + yPy}{N_p D}, \tag{S1.45}$$

where

$$xPx = \sum_{n=1}^{N} P^\top 1_n \|\mathbf{x}_n\|^2, \tag{S1.46}$$

$$yPy = \sum_{m=1}^{M} P1_m \|T\mathbf{Y}_m\|^2, \tag{S1.47}$$

$$\operatorname{tr}(PXY) = \sum_{m=1}^{M} (T\mathbf{Y}_m)^\top P X_m. \tag{S1.48}$$

A small variance floor prevents numerical collapse.

#### S1.8 Sparse k-nearest-neighbor E-step

The dense E-step evaluates all $MN$ source–target distances. For large atlas registrations, the implementation optionally switches to a sparse $k$-nearest-neighbor approximation after variance becomes sufficiently small. Each transformed source point queries a target k-d tree, and responsibilities are computed only for retained candidate pairs:

$$\widetilde{P}_{mn} \propto \exp\left[-\frac{\|\mathbf{x}_n - T\mathbf{Y}_m\|^2}{2\sigma^2}\right]. \tag{S1.49}$$

The same outlier constant is included in the denominator. Optional radius gating excludes candidates beyond a numerically safe variance-dependent radius. This approximation changes the responsibility sparsity pattern but not the M-step equations once $(P1, P^{\mathsf{T}}1, PX, N_p)$ are accumulated.

#### S1.9 Normalization and denormalization

In the benchmark, an outer preprocessing transform first expresses source and target coordinates in a target-scaled frame whose target bounding-box diagonal is 20. This step occurs outside the SSM-CPD class and is inverted before landmark errors are reported. The equations below describe the additional internal normalization performed by SSM-CPD.

When normalization is enabled, the target is centered and scaled:

$$\mathbf{X}' = \frac{\mathbf{X} - \mathbf{c}}{a}, \qquad \mathbf{c} = \frac{1}{N}\sum_{n=1}^{N}\mathbf{x}_n, \qquad a = \sqrt{\frac{1}{N}\sum_{n=1}^{N}\|\mathbf{x}_n - \mathbf{c}\|^2}. \tag{S1.50}$$

The source and basis use the same coordinate system:

$$\mathbf{Y}' = \frac{\mathbf{Y} - \mathbf{c}}{a}, \qquad \mathbf{U}' = \frac{\mathbf{U}}{a}. \tag{S1.51}$$

After convergence,

$$T\mathbf{Y} = aT\mathbf{Y}' + \mathbf{c}. \tag{S1.52}$$

If $T'(\mathbf{z}') = s\mathbf{R}\mathbf{z}' + \mathbf{t}'$, then the equivalent world-space transform is

$$T(\mathbf{z}) = s\mathbf{R}\mathbf{z} + \mathbf{t}_{\text{world}}, \qquad \mathbf{t}_{\text{world}} = -s\mathbf{R}\mathbf{c} + a\mathbf{t}' + \mathbf{c}. \tag{S1.53}$$

Under the implementation's row-vector convention, $\mathbf{t}_{\text{world}} = -s(\mathbf{c}\mathbf{R}^{\mathsf{T}}) + a\mathbf{t}' + \mathbf{c}$.

#### S1.10 Complete algorithm

1. Initialize $T\mathbf{Y} = \mathbf{Y}$, $\mathbf{R} = \mathbf{I}$, $s = 1$, $\mathbf{t} = \mathbf{0}$, $\mathbf{b} = \mathbf{0}$, and $\sigma^2$.

2. E-step: compute responsibilities $P_{mn}$ from the current transformed source and target points.

3. Accumulate $P1$, $P^{\mathsf{T}}1$, $PX$, and $N_p$.

4. Compute posterior target means $\bar{\mathbf{x}}_m = PX_m/P1_m$.

5. Apply the inverse similarity transform to obtain $\mathbf{i}_m = T^{-1}(\bar{\mathbf{x}}_m)$.
6. Solve
$$\left(\mathbf{U}^\mathsf{T}\mathbf{W}\mathbf{U} + \frac{\lambda_{\text{reg}}\sigma^2}{s^2}\mathbf{\Lambda}^{-1}\right)\mathbf{b} = \mathbf{U}^\mathsf{T}\mathbf{W}\mathbf{e}.$$
7. Form $\mathbf{Y}_b = \mathbf{Y} + \text{unvec}(\mathbf{U}\mathbf{b})$.
8. If enabled, update $(\mathbf{R}, s, \mathbf{t})$ by weighted Procrustes.
9. Set $T\mathbf{Y} = s\mathbf{R}\mathbf{Y}_b + \mathbf{t}$.
10. Update $\sigma^2$ from the weighted residuals.
11. Iterate until variance and coefficient changes meet the convergence tolerance or the maximum iteration count is reached.

#### S1.11 Limiting cases and interpretation

If $\lambda_{\text{reg}} \to 0$, the coefficient prior vanishes and coefficients are estimated by weighted least squares in the retained SSM subspace. The deformation remains restricted to the SSM span.

If $\lambda_{\text{reg}} \to \infty$, $\mathbf{b}$ is driven toward zero and the method approaches rigid or similarity registration of the undeformed atlas.

If $K = MD$ and $\mathbf{U}$ spans the full coordinate space, the SSM subspace no longer reduces dimensionality. In the implementation evaluated here, a truncated SSM restricts deformation to learned population variation.

When conventional CPD refinement follows SSM-CPD, the first stage supplies a biologically plausible coarse correspondence and the second permits additional local geometric adjustment.

#### S1.12 Implementation mapping

Implementation mapping between the mathematical derivation and software variables.

| Object | Meaning | Implementation |
|---|---|---|
| $\mathbf{X}$ | Target point cloud | `self.X` |
| $\mathbf{Y}$ | Source atlas correspondences or mean shape | `self.Y` |
| $\mathbf{U}$ | Flattened SSM basis | `self.U_flat` |
| $\mathbf{\Lambda}$ | SSM eigenvalue matrix | `self.L`, `self.invL` |
| $\mathbf{b}$ | SSM coefficient vector | `self.b` |
| $\mathbf{P}$ | Posterior responsibilities | `self.P` or implicit statistics |
| $P1$ | Responsibility row sums | `self.P1` |
| $P^{\mathsf{T}}1$ | Responsibility column sums | `self.Pt1` |
| $PX$ | Responsibility-weighted target sums | `self.PX` |
| $\bar{\mathbf{x}}$ | Posterior target means | `xbar` |
| $T^{-1}(\bar{\mathbf{x}})$ | Inverse-similarity target means | `ix` |
| $\mathbf{e}$ | Flattened inverse-frame residual | `E` |
| $\gamma$ | Prior scale $\lambda_{\text{reg}}\sigma^2/s^2$ | `gamma` |
| $\mathbf{A}$ | Coefficient normal-equation matrix | `system` |
| $\mathbf{U}^{\mathsf{T}}\mathbf{W}\mathbf{e}$ | Coefficient right-hand side | `rhs` |
| $\mathbf{Y}_b$ | SSM-deformed source | `Yb` |
| $T\mathbf{Y}$ | Transformed source points | `self.TY` |
| $\sigma^2$ | Residual variance | `self.sigma2` |

#### S1.13 Estimator summary and biological interpretation

The estimator performs MAP estimation of a CPD transformation whose non-rigid displacement is parameterized by an SSM:

$$T(\mathbf{Y}; \mathbf{b}, \mathbf{R}, s, \mathbf{t}) = s\mathbf{R}\left[\mathbf{Y} + \operatorname{unvec}(\mathbf{U}\mathbf{b})\right] + \mathbf{t}, \tag{S1.54}$$

with posterior correspondences estimated by the standard CPD E-step and coefficients estimated by Equation S1.28. The biological prior is explicit: deformation can follow observed population modes, and departures along low-variance modes are penalized more strongly than departures along high-variance modes.

### Supplementary Note S2: Integrated recovery workflow

#### S2.1 Scope and benchmark role

The integrated recovery workflow addresses cases in which a severely misaligned target lies outside the practical capture range of direct SSM-regularized CPD. It first searches a low-dimensional leave-one-out SSM for a target-compatible template, then estimates a feature-based global pose, and finally runs the same SSM-CPD and conventional-CPD refinement stages used in the primary benchmark. The workflow uses only the target surface and the training-fold SSM. Expert landmarks on the held-out target are never used for template selection, pose recovery, registration, or convergence decisions.

Although motivated as a rescue procedure, the severe-perturbation benchmark applied the complete sequence to every held-out target rather than selecting cases using their final landmark error. This prevents an outcome-dependent trigger from inflating the apparent recovery benefit. The complete sequence is interpreted as one estimator.

#### S2.2 Search template parameterization

Let $\bar{\mathbf{Y}} \in \mathbb{R}^{M\times 3}$ be the leave-one-out SSM mean, $\mathbf{u}_j \in \mathbb{R}^{3M}$ the $j$th unit-length PCA direction, and $\lambda_j$ its eigenvalue. The template-search implementation uses the first $k = 2$ modes and standardized search coordinates $b_j$:

$$\mathbf{Y}(\mathbf{b}) = \bar{\mathbf{Y}} + \mathrm{unvec}\left[\sum_{j=1}^{k} \mathbf{u}_j \sqrt{\lambda_j}\, b_j\right], \qquad -3 \le b_j \le 3. \tag{S2.1}$$

Thus, one unit of $b_j$ produces a one-standard-deviation displacement along mode $j$. This search parameterization is separate from the coefficient update derived in Supplementary Note S1.

For exact correspondence with the benchmark implementation, the template search additionally computes

$$\Omega(\mathbf{b}) = \sum_{j=1}^{k} \frac{b_j^2}{\lambda_j}. \tag{S2.2}$$

Because Equation S2.1 already expresses $b_j$ in standard- deviation units, Equation S2.2 should be understood as the implemented search regularizer, not as a re-expression of the SSM-CPD MAP prior in Supplementary Note S1.

#### S2.3 Feature-based template score

For a candidate $\mathbf{Y}(\mathbf{b})$, FPFH descriptors are computed from surface normals and used for RANSAC feature matching (Rusu *et al.*, 2009; Fischler & Bolles, 1981). Let $F(\mathbf{b})$ denote the RANSAC inlier fitness and let $R_{\mathrm{n}}(\mathbf{b})$ denote inlier RMSE divided by the bounding-box diagonal of the temporarily scaled target. Candidates maximize

$$J(\mathbf{b}) = F(\mathbf{b}) - 0.60 R_{\mathrm{n}}(\mathbf{b}) - w_{\mathrm{reg}}\Omega(\mathbf{b}), \tag{S2.3}$$

where

$$w_{\mathrm{reg}} = \min\left(\frac{0.10}{k}, \frac{0.25}{k(3)^2}\right) = \frac{1}{72} \qquad (k = 2). \tag{S2.4}$$

The score is evaluated only from target geometry and the training-fold SSM.

During template scoring, the target is temporarily rescaled so its bounding-box diagonal matches that of the SSM mean. With the template-search point-density parameter $p = 1.0$, the fine target voxel size is the scaled target diagonal divided by $25p$; the coarse voxel size is 1.5 times larger. Normal-search and FPFH-search radii are 2 and 5 voxel widths, respectively, at the fine scale and 1 and 2 voxel widths at the coarse scale. The RANSAC correspondence threshold is 3 voxel widths, the edge-length consistency threshold is 0.9, and confidence is 0.95. Coarse candidate point clouds are capped at 1,800 points; fine scoring uses the complete candidate cloud.

#### S2.4 Deterministic coefficient search

The coefficient search is deterministic (seed 0) and uses the following sequence.

1. Generate 192 scrambled Sobol low-discrepancy candidates in $[-3, 3]^2$ (Sobol', 1967).

2. Add heavy-tailed candidates generated from a two-dimensional Student $t$ distribution with two degrees of freedom, normalized by candidate RMS and assigned radii between 1.2 and 3.0; add axis-aligned candidates at ±3 and ±1.5 standard deviations.

3. Remove duplicates, standardize the candidate coordinates, embed the pool in two dimensions by singular-value decomposition, and use farthest-point sampling to retain a diverse beam of 24 candidates.

4. Coarsely score the beam with a nominal RANSAC budget of 600 iterations distributed across two restarts; retain up to 24 diverse candidates.

5. Fine-score the survivors with a nominal budget of 4,500 iterations distributed across four restarts; retain up to 12 diverse candidates.

6. Starting from the best candidate, run bounded Powell optimization (Powell, 1964) with at most 120 additional score evaluations, each using a nominal RANSAC budget of 30,000 iterations distributed across four restarts, and coefficient bounds $[-3, 3]^2$.

7. Re-evaluate the optimized candidate with a nominal final RANSAC budget of 150,000 iterations distributed across eight restarts and retain the best score encountered over all stages.

The resulting coefficient vector defines the optimized dense template through Equation S2.1. Sparse atlas landmarks are moved from the original atlas to this optimized template by thin-plate-spline interpolation with zero smoothing (Bookstein, 1989).

#### S2.5 Global pose recovery

The optimized template is next aligned to the target with a feature-based FPFH/RANSAC procedure followed by point-to-plane ICP (Rusu *et al.*, 2009; Fischler & Bolles, 1981; Besl & McKay, 1992; Rusinkiewicz & Levoy, 2001). Before descriptor matching, the optimized source is uniformly scaled so that its bounding-box diagonal matches the target diagonal; target coverage is fixed at 1.0. Source and target are independently normalized for feature calculation and then returned to their scaled coordinates. The descriptor voxel size is $1/(55p)$ in the normalized target frame, with $p = 1.0$; normal and FPFH radii are again 2 and 5 voxel widths, and the RANSAC distance threshold is 3 voxel widths.

The first RANSAC pass estimates a rigid transformation. Fitness is evaluated in both directions and averaged. If this symmetric fitness is below 0.99, up to 10 additional RANSAC passes permit uniform scale and the best result is retained. Each pass uses a maximum of 150,000 RANSAC iterations and confidence 0.95. The selected transformation is refined by point-to-plane ICP with a correspondence threshold of 0.4 voxel widths. Consequently, "rigid recovery" refers to the global pose-recovery stage, but the implementation can retain a uniform-scale correction when it improves the symmetric feature-match score.

#### S2.6 Non-rigid correspondence and landmark transfer

The pose-recovered optimized template initializes the SSM-CPD estimator described in Supplementary Note S1. The recovered source coordinates and consistently transformed shape modes are passed to the SSM-regularized estimator, which jointly re-estimates similarity and SSM coefficients using 95% SSM variance retention, $\lambda_{\text{reg}} = 0.01$, $w = 0.1$, $k = 10$, a maximum of 250 iterations, tolerance $10^{-6}$, and joint scale, rotation, and translation. The resulting dense correspondences are then refined with conventional deformable

CPD (Myronenko & Song, 2010), using $\alpha = 2$, $\beta = 2$, $w = 0.1$, at most 80 low-rank kernel modes, $k = 10$, a maximum of 250 iterations, and tolerance $10^{-6}$. Finally, sparse landmarks are transferred by a zero-smoothing thin-plate spline from the pre-registration dense source to the final dense correspondences (Bookstein, 1989).

The complete severe-perturbation sequence is therefore

$$\begin{aligned}\text{SSM template search} &\longrightarrow \text{FPFH/RANSAC + ICP pose recovery}\\ &\longrightarrow \text{SSM-CPD} \longrightarrow \text{CPD refinement} \longrightarrow \text{sparse landmark transfer.}\end{aligned} \tag{S2.5}$$

All stochastic components use fixed seeds, and all hyperparameters are shared across datasets and specimens. The benchmark caches deterministic preprocessing products for efficiency, but cache use does not change the estimated transformation or landmark predictions.

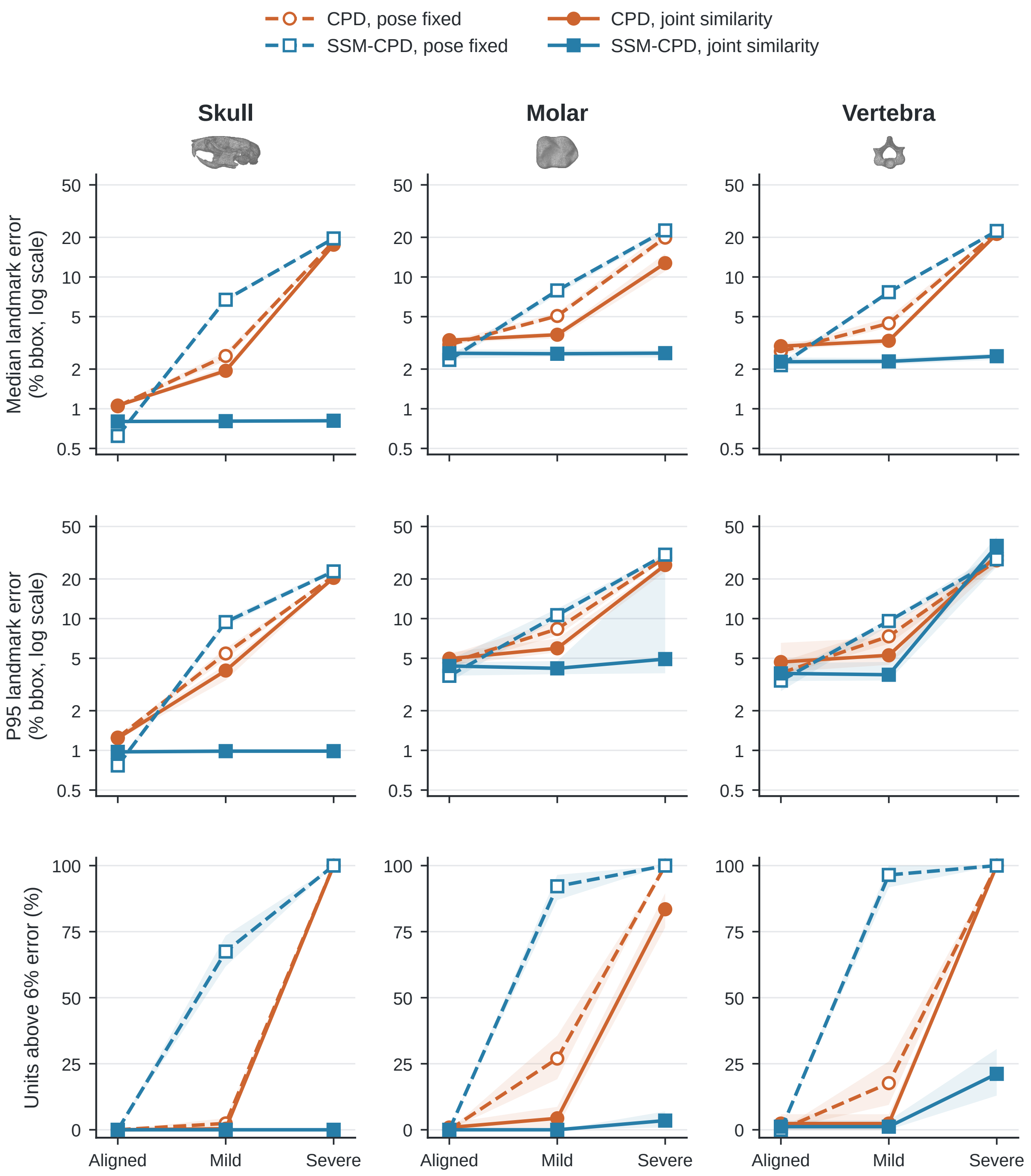

**Figure S1:** Pose-by-formulation attribution across initialization regimes. Rows report median landmark error, P95 landmark error, and the fraction of independent units above the 6% failure threshold, computed over independent units (specimens for skulls and molars; individual-level medians for vertebrae). Dashed lines with open markers denote pose-fixed conditions; solid lines with filled markers denote joint similarity optimization. Bands show 95% biological-unit bootstrap intervals based on 2,000 resamples.

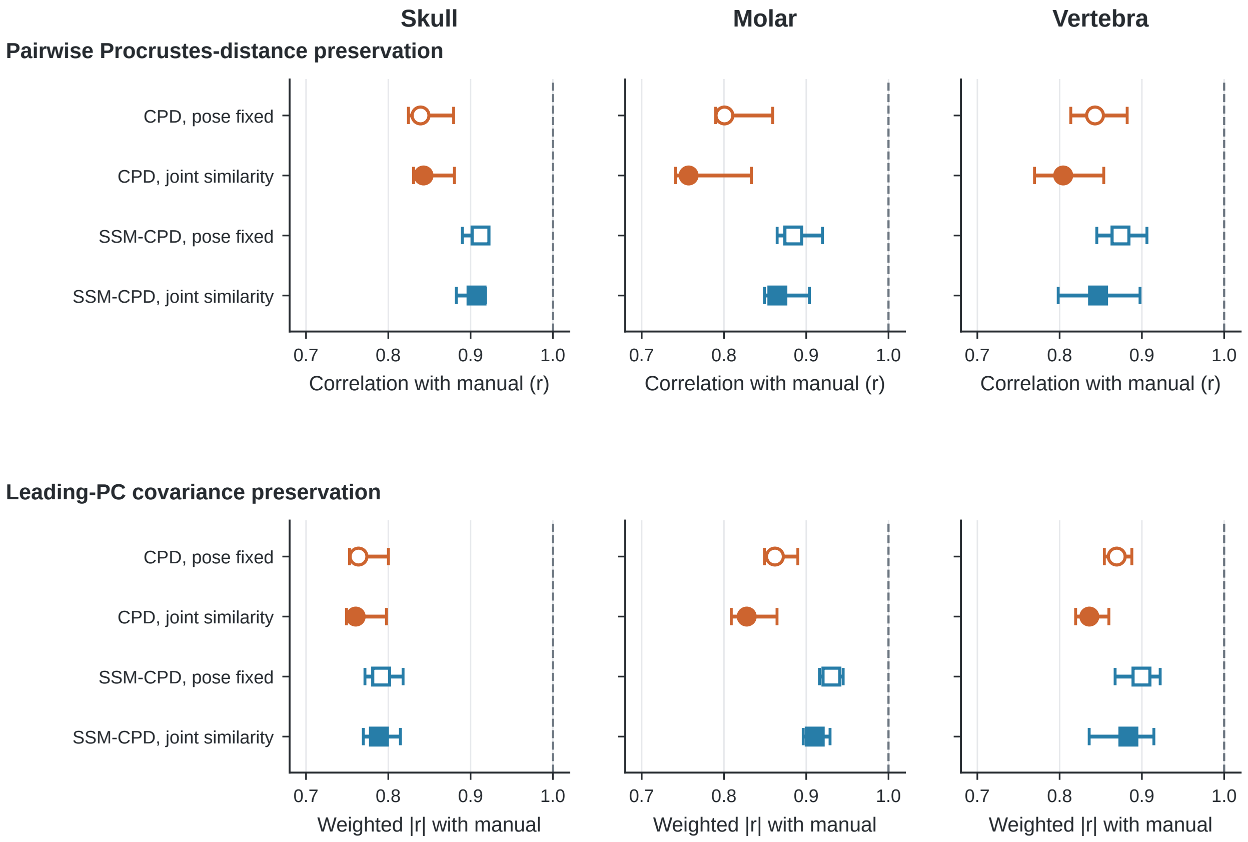

**Figure S2:** Aligned-regime shape preservation in the four-cell attribution design. Top: pairwise Procrustes-distance preservation. Bottom: variance-weighted absolute PC-score correlation across the manual principal components explaining 80% of manual shape variance. Open markers denote pose-fixed conditions and filled markers joint similarity optimization. Points are observed correlations and intervals are 95% bootstrap intervals based on 2,000 resamples of biological independent units.

**Table S1:** Complete paired effects for the 2 × 2 pose-by-formulation attribution analysis.

| Dataset | Regime | Effect | $n$ | Median Δ | 95% CI | Win rate |
|---|---|---|---|---|---|---|
| Skull | Aligned | Pose effect within CPD | 249 | -0.01 | [-0.01, 0.00] | 42.2% |
| Skull | Aligned | Pose effect within SSM-CPD | 249 | -0.15 | [-0.18, -0.14] | 0.4% |
| Skull | Aligned | Formulation effect, pose fixed | 249 | 0.41 | [0.39, 0.43] | 100.0% |
| Skull | Aligned | Formulation effect, joint similarity | 249 | 0.26 | [0.25, 0.27] | 99.2% |
| Skull | Aligned | Pose × formulation interaction | 249 | 0.14 | [0.12, 0.16] | 94.8% |
| Skull | Mild | Pose effect within CPD | 249 | 0.44 | [0.36, 0.54] | 92.8% |
| Skull | Mild | Pose effect within SSM-CPD | 249 | 5.95 | [5.72, 6.23] | 100.0% |
| Skull | Mild | Formulation effect, pose fixed | 249 | -3.89 | [-4.05, -3.75] | 0.0% |
| Skull | Mild | Formulation effect, joint similarity | 249 | 1.16 | [1.05, 1.26] | 100.0% |
| Skull | Mild | Pose × formulation interaction | 249 | -5.28 | [-5.48, -5.10] | 0.0% |
| Skull | Severe | Pose effect within CPD | 249 | 1.11 | [0.93, 1.27] | 88.4% |
| Skull | Severe | Pose effect within SSM-CPD | 249 | 18.79 | [18.40, 19.06] | 100.0% |
| Skull | Severe | Formulation effect, pose fixed | 249 | -1.36 | [-1.55, -1.01] | 19.3% |
| Skull | Severe | Formulation effect, joint similarity | 249 | 16.72 | [16.34, 16.91] | 100.0% |
| Skull | Severe | Pose × formulation interaction | 249 | -17.67 | [-17.96, -17.29] | 0.0% |
| Molar | Aligned | Pose effect within CPD | 115 | -0.23 | [-0.36, -0.12] | 30.4% |
| Molar | Aligned | Pose effect within SSM-CPD | 115 | -0.18 | [-0.29, -0.10] | 34.8% |
| Molar | Aligned | Formulation effect, pose fixed | 115 | 0.59 | [0.45, 0.78] | 81.7% |
| Molar | Aligned | Formulation effect, joint similarity | 115 | 0.72 | [0.63, 0.89] | 84.3% |
| Molar | Aligned | Pose × formulation interaction | 115 | -0.03 | [-0.15, 0.10] | 49.6% |
| Molar | Mild | Pose effect within CPD | 115 | 1.19 | [1.01, 1.56] | 90.4% |
| Molar | Mild | Pose effect within SSM-CPD | 115 | 4.97 | [4.48, 5.46] | 100.0% |
| Molar | Mild | Formulation effect, pose fixed | 115 | -2.62 | [-2.89, -2.45] | 1.7% |
| Molar | Mild | Formulation effect, joint similarity | 115 | 0.98 | [0.69, 1.22] | 87.8% |
| Molar | Mild | Pose × formulation interaction | 115 | -3.55 | [-3.96, -3.31] | 0.0% |
| Molar | Severe | Pose effect within CPD | 115 | 7.65 | [6.19, 8.53] | 97.4% |
| Molar | Severe | Pose effect within SSM-CPD | 115 | 19.60 | [18.36, 20.35] | 97.4% |
| Molar | Severe | Formulation effect, pose fixed | 115 | -2.07 | [-2.35, -1.59] | 13.9% |
| Molar | Severe | Formulation effect, joint similarity | 115 | 9.52 | [7.65, 11.13] | 97.4% |
| Molar | Severe | Pose × formulation interaction | 115 | -11.22 | [-13.43, -10.08] | 2.6% |
| Vertebra | Aligned | Pose effect within CPD | 85 | -0.28 | [-0.34, -0.24] | 18.8% |
| Vertebra | Aligned | Pose effect within SSM-CPD | 85 | -0.18 | [-0.23, -0.12] | 22.4% |
| Vertebra | Aligned | Formulation effect, pose fixed | 85 | 0.57 | [0.49, 0.66] | 85.9% |
| Vertebra | Aligned | Formulation effect, joint similarity | 85 | 0.60 | [0.48, 0.73] | 88.2% |
| Vertebra | Aligned | Pose × formulation interaction | 85 | -0.14 | [-0.25, 0.02] | 42.4% |
| Vertebra | Mild | Pose effect within CPD | 85 | 1.17 | [0.80, 1.43] | 95.3% |
| Vertebra | Mild | Pose effect within SSM-CPD | 85 | 5.20 | [5.09, 5.48] | 98.8% |
| Vertebra | Mild | Formulation effect, pose fixed | 85 | -2.95 | [-3.20, -2.58] | 0.0% |
| Vertebra | Mild | Formulation effect, joint similarity | 85 | 0.94 | [0.81, 1.08] | 94.1% |
| Vertebra | Mild | Pose × formulation interaction | 85 | -4.02 | [-4.30, -3.69] | 1.2% |
| Vertebra | Severe | Pose effect within CPD | 85 | 0.70 | [0.11, 1.58] | 62.4% |
| Vertebra | Severe | Pose effect within SSM-CPD | 85 | 18.61 | [17.87, 19.64] | 85.9% |
| Vertebra | Severe | Formulation effect, pose fixed | 85 | -0.84 | [-1.26, -0.34] | 31.8% |
| Vertebra | Severe | Formulation effect, joint similarity | 85 | 16.09 | [15.30, 17.43] | 83.5% |
| Vertebra | Severe | Pose × formulation interaction | 85 | -17.01 | [-18.76, -15.67] | 14.1% |

*Note.* Positive reductions favor the second condition in each contrast. Confidence intervals are percentile bootstrap intervals at the biological-independent-unit level.

**Table S2:** Four-cell independent-unit summaries for the pose-by-formulation attribution analysis.

| Dataset | Regime | Method | $n$ | Median (95% CI) | P90 (95% CI) | P95 (95% CI) | Failure > 6% (95% CI) |
|---|---|---|---|---|---|---|---|
| Skull | Aligned | CPD · pose fixed | 249 | 1.05 [1.02, 1.07] | 1.19 [1.17, 1.23] | 1.25 [1.21, 1.29] | 0.0% [0.0%, 0.0%] |
| Skull | Aligned | CPD · joint similarity | 249 | 1.06 [1.04, 1.08] | 1.20 [1.17, 1.22] | 1.24 [1.20, 1.30] | 0.0% [0.0%, 0.0%] |
| Skull | Aligned | SSM-CPD · pose fixed | 249 | 0.62 [0.61, 0.63] | 0.72 [0.71, 0.75] | 0.77 [0.73, 0.81] | 0.0% [0.0%, 0.0%] |
| Skull | Aligned | SSM-CPD · joint similarity | 249 | 0.80 [0.79, 0.82] | 0.93 [0.91, 0.95] | 0.97 [0.94, 1.00] | 0.0% [0.0%, 0.0%] |
| Skull | Mild | CPD · pose fixed | 249 | 2.51 [2.32, 2.72] | 4.73 [4.19, 5.14] | 5.44 [4.94, 5.95] | 2.4% [0.8%, 4.4%] |
| Skull | Mild | CPD · joint similarity | 249 | 1.94 [1.83, 2.09] | 3.32 [3.12, 3.78] | 4.04 [3.40, 4.49] | 0.4% [0.0%, 1.2%] |
| Skull | Mild | SSM-CPD · pose fixed | 249 | 6.72 [6.48, 7.01] | 8.49 [8.22, 9.10] | 9.42 [8.77, 9.98] | 67.5% [61.8%, 73.5%] |
| Skull | Mild | SSM-CPD · joint similarity | 249 | 0.81 [0.79, 0.82] | 0.93 [0.91, 0.96] | 0.99 [0.95, 1.02] | 0.0% [0.0%, 0.0%] |
| Skull | Severe | CPD · pose fixed | 249 | 18.56 [18.25, 18.84] | 20.67 [20.27, 20.88] | 21.01 [20.82, 21.46] | 100.0% [100.0%, 100.0%] |
| Skull | Severe | CPD · joint similarity | 249 | 17.57 [17.14, 17.76] | 19.75 [19.32, 20.14] | 20.38 [19.89, 20.61] | 100.0% [100.0%, 100.0%] |
| Skull | Severe | SSM-CPD · pose fixed | 249 | 19.61 [19.25, 19.86] | 22.08 [21.71, 22.39] | 22.82 [22.35, 23.97] | 100.0% [100.0%, 100.0%] |
| Skull | Severe | SSM-CPD · joint similarity | 249 | 0.81 [0.79, 0.83] | 0.95 [0.93, 0.97] | 0.99 [0.96, 1.03] | 0.0% [0.0%, 0.0%] |
| Molar | Aligned | CPD · pose fixed | 115 | 3.06 [2.93, 3.27] | 4.28 [3.88, 4.53] | 4.61 [4.27, 5.33] | 0.0% [0.0%, 0.0%] |
| Molar | Aligned | CPD · joint similarity | 115 | 3.31 [3.17, 3.43] | 4.41 [4.12, 4.86] | 4.97 [4.38, 5.53] | 0.9% [0.0%, 2.6%] |
| Molar | Aligned | SSM-CPD · pose fixed | 115 | 2.35 [2.20, 2.51] | 3.31 [3.10, 3.63] | 3.68 [3.30, 4.86] | 0.0% [0.0%, 0.0%] |
| Molar | Aligned | SSM-CPD · joint similarity | 115 | 2.64 [2.40, 2.81] | 3.69 [3.40, 4.25] | 4.37 [3.69, 4.66] | 0.0% [0.0%, 0.0%] |
| Molar | Mild | CPD · pose fixed | 115 | 5.06 [4.90, 5.38] | 7.41 [6.41, 8.28] | 8.35 [7.41, 9.63] | 27.0% [19.1%, 35.7%] |
| Molar | Mild | CPD · joint similarity | 115 | 3.65 [3.41, 3.81] | 5.54 [4.94, 5.95] | 5.95 [5.55, 7.06] | 4.3% [0.9%, 8.7%] |
| Molar | Mild | SSM-CPD · pose fixed | 115 | 7.91 [7.38, 8.20] | 9.98 [9.57, 10.52] | 10.63 [9.96, 11.89] | 92.2% [87.0%, 96.5%] |
| Molar | Mild | SSM-CPD · joint similarity | 115 | 2.61 [2.49, 2.82] | 3.84 [3.50, 4.15] | 4.19 [3.78, 4.76] | 0.0% [0.0%, 0.0%] |
| Molar | Severe | CPD · pose fixed | 115 | 19.89 [18.61, 22.39] | 28.03 [26.83, 28.61] | 29.03 [28.04, 30.92] | 100.0% [100.0%, 100.0%] |
| Molar | Severe | CPD · joint similarity | 115 | 12.73 [11.03, 15.07] | 22.73 [20.70, 25.34] | 25.50 [22.65, 26.32] | 83.5% [76.5%, 89.6%] |
| Molar | Severe | SSM-CPD · pose fixed | 115 | 22.57 [21.34, 23.62] | 29.30 [27.80, 30.53] | 30.57 [29.38, 31.02] | 100.0% [100.0%, 100.0%] |
| Molar | Severe | SSM-CPD · joint similarity | 115 | 2.64 [2.44, 2.79] | 3.85 [3.38, 4.78] | 4.93 [3.85, 27.75] | 3.5% [0.9%, 7.0%] |
| Vertebra | Aligned | CPD · pose fixed | 85 | 2.72 [2.55, 2.88] | 3.78 [3.42, 3.87] | 3.86 [3.72, 4.69] | 0.0% [0.0%, 0.0%] |
| Vertebra | Aligned | CPD · joint similarity | 85 | 2.99 [2.84, 3.22] | 4.00 [3.76, 4.72] | 4.68 [3.94, 6.55] | 2.4% [0.0%, 5.9%] |
| Vertebra | Aligned | SSM-CPD · pose fixed | 85 | 2.14 [2.02, 2.25] | 2.93 [2.64, 3.39] | 3.38 [2.81, 3.68] | 0.0% [0.0%, 0.0%] |
| Vertebra | Aligned | SSM-CPD · joint similarity | 85 | 2.27 [2.15, 2.45] | 3.40 [2.92, 3.87] | 3.84 [3.36, 4.59] | 1.2% [0.0%, 3.5%] |
| Vertebra | Mild | CPD · pose fixed | 85 | 4.45 [4.16, 4.95] | 6.63 [5.98, 7.39] | 7.34 [6.43, 8.64] | 17.6% [9.4%, 25.9%] |
| Vertebra | Mild | CPD · joint similarity | 85 | 3.28 [3.08, 3.43] | 4.52 [4.25, 5.24] | 5.27 [4.46, 7.18] | 2.4% [0.0%, 5.9%] |
| Vertebra | Mild | SSM-CPD · pose fixed | 85 | 7.67 [7.33, 7.82] | 8.98 [8.50, 9.58] | 9.59 [8.81, 9.94] | 96.5% [91.8%, 100.0%] |
| Vertebra | Mild | SSM-CPD · joint similarity | 85 | 2.29 [2.17, 2.43] | 3.49 [2.82, 3.75] | 3.75 [3.39, 4.69] | 1.2% [0.0%, 3.5%] |
| Vertebra | Severe | CPD · pose fixed | 85 | 21.90 [21.45, 22.51] | 25.36 [24.21, 27.86] | 27.65 [25.11, 31.44] | 100.0% [100.0%, 100.0%] |
| Vertebra | Severe | CPD · joint similarity | 85 | 21.21 [20.37, 22.20] | 25.97 [24.24, 30.63] | 30.07 [25.34, 32.04] | 100.0% [100.0%, 100.0%] |
| Vertebra | Severe | SSM-CPD · pose fixed | 85 | 22.36 [21.71, 23.04] | 27.21 [25.34, 28.22] | 28.20 [26.69, 31.68] | 100.0% [100.0%, 100.0%] |
| Vertebra | Severe | SSM-CPD · joint similarity | 85 | 2.51 [2.35, 2.63] | 27.79 [17.15, 35.68] | 35.64 [24.68, 41.97] | 21.2% [12.9%, 30.6%] |

*Note.* Error is the median sparse-landmark error as a percentage of the subsampled-target bounding-box diagonal. Summaries are computed over independent units: specimens for skulls and molars and the 85 individual-level medians for vertebrae; intervals resample the same units.

**Table S3:** Shape preservation across the aligned four-cell attribution design.

| Dataset | Method | $n$ spec./units | Pairwise $r$ (95% CI) | PC1 $r$ (95% CI) | PC2 $r$ (95% CI) | PC80 trace (95% CI) |
|---|---|---|---|---|---|---|
| Skull | CPD · pose fixed | 249/249 | 0.84 [0.82, 0.88] | 0.96 [0.95, 0.98] | 0.93 [0.89, 0.97] | 0.56 [0.49, 0.54] |
| Skull | CPD · joint similarity | 249/249 | 0.84 [0.83, 0.88] | 0.96 [0.95, 0.98] | 0.93 [0.89, 0.96] | 0.56 [0.49, 0.53] |
| Skull | SSM-CPD · pose fixed | 249/249 | 0.91 [0.89, 0.92] | 0.98 [0.97, 0.98] | 0.97 [0.95, 0.97] | 0.39 [0.36, 0.40] |
| Skull | SSM-CPD · joint similarity | 249/249 | 0.91 [0.88, 0.92] | 0.98 [0.97, 0.99] | 0.97 [0.95, 0.98] | 0.35 [0.33, 0.37] |
| Molar | CPD · pose fixed | 115/115 | 0.80 [0.79, 0.86] | 0.95 [0.93, 0.96] | 0.91 [0.81, 0.95] | 0.25 [0.22, 0.27] |
| Molar | CPD · joint similarity | 115/115 | 0.76 [0.74, 0.83] | 0.93 [0.91, 0.96] | 0.92 [0.71, 0.95] | 0.23 [0.19, 0.24] |
| Molar | SSM-CPD · pose fixed | 115/115 | 0.88 [0.86, 0.92] | 0.99 [0.98, 0.99] | 0.96 [0.91, 0.97] | 0.74 [0.68, 0.76] |
| Molar | SSM-CPD · joint similarity | 115/115 | 0.86 [0.85, 0.90] | 0.98 [0.97, 0.99] | 0.95 [0.87, 0.97] | 0.71 [0.65, 0.74] |
| Vertebra | CPD · pose fixed | 254/85 | 0.84 [0.81, 0.88] | 0.96 [0.94, 0.97] | 0.89 [0.82, 0.93] | 0.38 [0.34, 0.41] |
| Vertebra | CPD · joint similarity | 254/85 | 0.80 [0.77, 0.85] | 0.94 [0.92, 0.96] | 0.85 [0.78, 0.91] | 0.34 [0.30, 0.37] |
| Vertebra | SSM-CPD · pose fixed | 254/85 | 0.87 [0.85, 0.91] | 0.98 [0.97, 0.99] | 0.93 [0.86, 0.96] | 0.74 [0.68, 0.77] |
| Vertebra | SSM-CPD · joint similarity | 254/85 | 0.85 [0.80, 0.90] | 0.96 [0.91, 0.99] | 0.93 [0.88, 0.96] | 0.75 [0.68, 0.79] |

*Note.* Intervals are percentile bootstrap intervals from 2,000 resamples. Vertebral resampling is clustered by biological individual. The GPA and manual PCA basis are refitted within each resample; percentile intervals summarize that refitted bootstrap distribution and are not mathematically constrained to contain the full-sample estimate. This explains the two skull CPD trace-ratio rows whose full-sample estimates are 0.56 and whose percentile intervals are [0.49, 0.54] and [0.49, 0.53].

**Table S4:** Highest-error cases and their response to integrated recovery.

| Dataset | Specimen | CPD + joint similarity | SSM-CPD + refinement | Integrated recovery | Recovery Δ | Recovery < 6% |
|---|---|---|---|---|---|---|
| Skull | `1255` | 22.06 | 1.05 | 1.03 | 0.02 | Yes |
| Skull | `889` | 22.06 | 0.92 | 0.92 | 0.00 | Yes |
| Skull | `1523` | 21.33 | 0.96 | 0.94 | 0.02 | Yes |
| Skull | `473` | 21.21 | 0.72 | 0.79 | -0.06 | Yes |
| Skull | `485` | 21.10 | 0.91 | 0.89 | 0.02 | Yes |
| Skull | `407` | 20.97 | 1.04 | 0.95 | 0.08 | Yes |
| Skull | `1074` | 20.91 | 1.07 | 1.09 | -0.02 | Yes |
| Skull | `1511` | 20.61 | 0.79 | 0.82 | -0.03 | Yes |
| Molar | `i14_sas_align` | 19.77 | 3.27 | 52.20 | -48.92 | No |
| Molar | `w02_sas_align` | 8.50 | 2.88 | 47.76 | -44.88 | No |
| Molar | `u16_sas_align` | 15.56 | 4.25 | 33.49 | -29.24 | No |
| Molar | `u20_sas_align` | 25.39 | 30.49 | 3.94 | 26.55 | Yes |
| Molar | `x21_sas_align` | 25.88 | 29.97 | 1.91 | 28.06 | Yes |
| Molar | `j11_sas_align` | 26.71 | 29.45 | 1.57 | 27.88 | Yes |
| Molar | `i26_sas_align` | 27.71 | 2.38 | 2.62 | -0.25 | Yes |
| Molar | `q19_sas_align` | 26.32 | 1.59 | 1.95 | -0.37 | Yes |
| Vertebra | `gerrhosauridae_tetradactylus_tetradactylus_ummz_08-t1` | 39.28 | 47.35 | 2.22 | 45.13 | Yes |
| Vertebra | `cordylidae_ouroborus_cataphractus_amnh_01-c3` | 32.26 | 46.72 | 2.77 | 43.96 | Yes |
| Vertebra | `gerrhosauridae_tetradactylus_tetradactylus_ummz_15-l1` | 37.10 | 45.65 | 2.05 | 43.60 | Yes |
| Vertebra | `gerrhosauridae_cordylosaurus_subtessellatus_mcz19331_07-t1` | 15.56 | 45.55 | 3.52 | 42.03 | Yes |
| Vertebra | `varanidae_varanus-prasinus-uf71411_15-l1` | 32.99 | 45.30 | 2.41 | 42.89 | Yes |
| Vertebra | `cordylidae_platysaurus_capensis_cas203515_01-c3` | 34.27 | 44.85 | 3.63 | 41.22 | Yes |
| Vertebra | `cordylidae_ninurta_coerleo_ni_15-l1` | 32.84 | 44.38 | 2.41 | 41.97 | Yes |
| Vertebra | `cordylidae_pseudocordylus_langi_ps-langi_21-l1` | 31.02 | 44.13 | 2.26 | 41.88 | Yes |

*Note.* Cases are the eight specimens per dataset with the largest severe-perturbation error under any of the three listed methods. Values are landmark error as a percentage of the subsampled-target bounding-box diagonal. Recovery Δ is SSM-CPD + refinement minus integrated recovery; negative values indicate that recovery increased error. Recovery < 6% indicates whether the integrated-recovery error is below the 6% failure threshold.

**Table S5:** Manual landmark quality-control summary.

| Dataset | *n* | LM | Unique | Duplicates | SD min | SD median | SD max | GPA rank | PCs80 |
|---|---|---|---|---|---|---|---|---|---|
| Skull | 249 | 79 | 249 | 0 | 0.0034 | 0.0047 | 0.0067 | 11.86 | 28 |
| Molar | 115 | 16 | 115 | 0 | 0.0310 | 0.0465 | 0.0803 | 6.86 | 9 |
| Vertebra | 254 | 28 | 254 | 0 | 0.0239 | 0.0359 | 0.0669 | 5.67 | 8 |

LM, sparse landmarks; SD, landmark coordinate standard deviation after GPA; GPA rank, effective rank of the manual GPA configuration; PCs80, number of principal components explaining 80% of manual shape variance.
